# Molecular architecture of synaptic vesicles

**DOI:** 10.1101/2024.04.11.588828

**Authors:** Uljana Kravcenko, Max Ruwolt, Jana Kroll, Artsemi Yushkevich, Martina Zenkner, Julia Ruta, Rowaa Lotfy, Erich E. Wanker, Christian Rosenmund, Fan Liu, Mikhail Kudryashev

## Abstract

Synaptic vesicles (SVs) store and transport neurotransmitters to the presynaptic active zone for release by exocytosis. After release, SV proteins and excess membrane are recycled via endocytosis, and new SVs are formed in a clathrin-dependent manner. This process maintains the morphology and complex molecular composition of SVs through multiple recycling rounds. Previous studies explored the molecular composition of SVs through proteomic analysis and fluorescent microscopy, proposing a model for an average SV^1,2^. However, the structural heterogeneity and molecular architecture of individual SVs are not well described. Here we used cryo-electron tomography to visualize morphological and molecular details of SVs isolated from mouse brains and inside cultured neurons. We describe several classes of small proteins on the SV surface and long proteinaceous densities inside SVs. We identified V-ATPases, determined a structure using subtomogram average, and showed them forming a complex with the membrane-embedded protein synaptophysin. Our bioluminescence assay revealed pairwise interactions between VAMP2 and synaptophysin and V-ATPase Voe1 domains. Interestingly, V-ATPases were randomly distributed on the surface of SVs irrespective of vesicle sizes. A subpopulation of isolated vesicles and vesicles inside neurons contained a partially assembled clathrin coat with a soccer-ball symmetry. We observed a V-ATPase under clathrin cage in several isolated clathrin-coated vesicles. Additionally, from isolated SV preparations and within hippocampal neurons we identified clathrin baskets without vesicles. We determined their preferential location in proximity to the cell membrane. Our analysis advances the understanding of individual SVs’ diversity and their molecular architecture.

## Introduction

Synaptic vesicles (SVs) display the fundamental elements of synaptic transmission and thus of neuronal communication. Previous studies demonstrated that the surface of SVs is crowded with more than 40 different proteins involved in neurotransmitter (NT) release and SV recycling^1,3,4^. The most abundant proteins are SNAREs (VAMPs, syntaxins), calcium sensors (synaptotagmins), endocytosis-related proteins (dynamins), small GTPases (Rabs), and other trafficking and membrane proteins (CSP)^1,5^. While the molecular composition of SVs has been well-studied using mass spectrometry^6,7^ and fluorescence microscopy^2,4^, the architecture of individual SVs has not been analyzed at molecular resolution.

Neuronal depolarization induces the fusion of NT-filled SVs with the presynaptic active zone membrane, leading to the release of their contents into the synaptic cleft and signal propagation to the postsynapse^8^. During subsequent endocytosis, SV proteins and excessive membranes are internalized and new SVs are reformed. To preserve the morphological and molecular organization of SVs throughout several rounds of SV fusion and recycling, these processes must be tightly controlled^9^. To couple exo- and endocytosis and to organize the proper reformation of SVs, highly abundant SV proteins VAMP2 and synaptophysin (Syp) interact with the clathrin adaptor AP180 and the GTPase dynamin at the periactive zone, respectively^10,11^. While the endocytic internalization of membranes can be clathrin-independent, e.g. via ultrafast endocytosis or activity-dependent bulk endocytosis, the reformation of new SVs is supposed to be clathrin-dependent^12^. It is likely that the organization of clathrin triskelia in pentamers and hexamers is required to produce membrane curvature and lead to the formation of vesicles with a defined size^13,14^. Clathrin-coated vesicles (CCVs) were shown to have several symmetry types defined mostly by the number and arrangement of pentagonal or hexagonal faces, which are organized by the adaptor proteins^15^. Interestingly, also non-vesicle-carrying clathrin cages (baskets) were observed to be formed *in vitro* and hypothesized to be assembled spontaneously from monomers during the isolation procedure^16,17^.

To reform fusion-competent SVs, CCVs must be uncoated, their lumen needs to be acidified via V-ATPases and NT must be loaded via transporters like VGLUT, VMAT, VAChT, and others. Short before SV fusion, the extravesicular V1 region of the V-ATPase, required for ATP hydrolysis, is dissociated from the membrane-embedded Vo region^18^. After SV fusion, the V1 region is recruited back to the SV, likely via rabconnectin-3^19^. Although immunoblot experiments have suggested that CCVs contain both Vo and V1 regions^20^, it is unclear whether the two domains are already assembled on CCVs. In contrast to the V1 region, the Vo region is supposed to remain integrated into the membrane during fusion and recycled during endocytosis. Considering that each SV has on average 1.4 V-ATPases^1^, it is conceivable that the tight control of the Vo region recycling and the V1 region recruitment are crucial for the proper timing of fast SV refilling, particularly during sustained NT release.

In the present study, we performed cryo-electron tomography (cryo-ET) on neuronal SVs and CCVs within cultured hippocampal mouse neurons *in situ* and isolated vesicles from mouse brains *ex vivo* to characterize their molecular architecture. This enabled us to morphologically categorize different types of proteins visible at the SV surface. Beyond that, subtomogram averaging (StA) revealed a structure of the V-ATPase interacting with a small transmembrane protein that we identified as Syp. We observed assembled V-ATPases under the cage of CCVs, and characterized partially (un)coated CCVs both *in situ* and in *ex vivo* preparations. Interestingly, we detected empty clathrin cages, performed StA on them, and characterized their size, type of symmetry, and preferred cellular localization.

## Results

### Cryo-ET of SVs from primary hippocampal neuron cultures *in situ* and isolated from mouse brains

We first aimed to visualize the composition of SVs *in situ*. For this, we imaged presynaptic areas of mouse hippocampal neurons cultured on electron microscopy grids (DIV 17) by cryo-ET (Fig.1 *A*). We recorded 88 tomograms containing 56 putative presynaptic terminals (Fig.1 *B*). As a second independent approach, we extracted vesicles from mouse brain tissue (hippocampus, cortex, and cerebellum) (Fig. 1 *C*). After confirming successful vesicle purification via mass-spectrometry (*SI*, Table S1), we recorded 719 tomograms containing vesicles purified from mouse brains (see Methods). These tomograms of isolated vesicles provided a higher signal-to-noise ratio and allowed visualization of the molecules on their surface in more detail (Fig. 1 *D*). To preserve the molecular architecture of vesicles as well as possible, we chose a gentle purification protocol. As a consequence, our vesicle preparation also contained fractions of other classes of vesicles like CCVs. In this manuscript, we identify SVs as vesicles containing at least one V-ATPase.

**Fig. 1.**
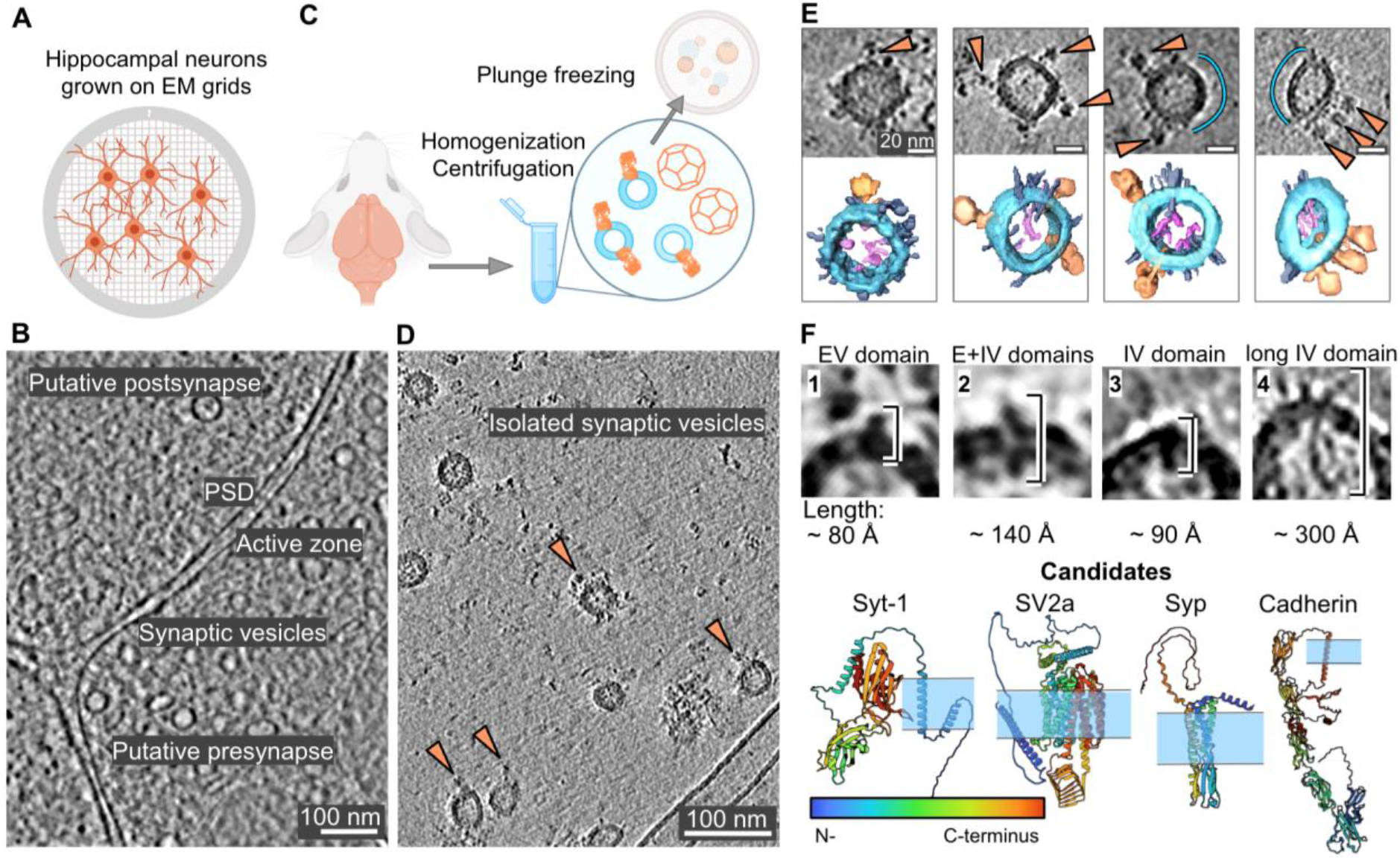
Visualization of individual SVs and proteins on their surface. A) Schematic representation of primary hippocampal neurons grown on grids. B) A slice through a cryo-ET volume of a putative presynapse *in situ* containing individual SVs. C) Schematic representation of SV isolation procedure. D) A slice through a cryo-ET volume of isolated SVs. The tomogram for this image was CTF deconvoluted using Isonet^21^. E) Upper row: isolated individual SVs, with proteins on their surface. Blue curves indicate SV surfaces without detectable proteins. Lower row: segmentations showing V-ATPase densities (orange), intravesicular protrusions (pink), and small membrane proteins (gray) on the surface of SVs (blue). F) Protein densities are grouped into classes based on their shapes in the tomograms of isolated SVs. Predictions of structures of potential candidates for observed densities from the Alphafold2^22^ database and a scheme of their membrane localization(the membrane is schematically shown in blue). N to C termini for framed AlphaFold-predicted proteins are shown from blue to red. V-ATPase PDB: 6wm2^23^. Scale bars: B), D) 100 nm, E) 20 nm. V-ATPases are shown with orange arrows inD) and E).

### Molecular landscape of synaptic vesicle membrane

The overall size and morphology of SVs *in situ* and of isolated SVs were comparable (Fig. 1, *SI Appendix, Fig. S1*). Characteristic protein densities particularly on the surface of SVs but also within the SV lumen were visible in both preparations (Fig. 1 *B, D*, *SI Appendix, Fig. S2*). Since the resolution of isolated SVs was higher than that of SVs *in situ* due to the lower thickness of vitreous ice, we focused on isolated SVs for protein identification and characterization. Interestingly, we could observe that while most of the surface of SVs was covered by protein densities, some areas did not have apparent proteins (Fig. 1 *E*, empty surface is shown with blue segments). We next classified the apparent protein densities into four structural classes, according to their size and membrane topology: (1) small extra-vesicular (EV) domain; (2) extra-and-intravesicular (E+IV) domains; (3) small intravesicular (IV) domain and (4) long IV domain (Fig. 1 *F*).

We aimed to quantify the abundance of proteins from different classes per SV. For this, five independent expert annotators who did not know the previously reported proteomic composition^1,6^, inspected 90 SVs to identify protein-like densities and assign each observed density to one of the four suggested structural classes. To estimate the abundance of proteins from outlined classes, the mean value of expert counts per each structural class was calculated for each SV individually, followed by averaging the obtained mean counts per class over all SVs. Quantified occurrences of smaller proteins per SV were statistically unreliable due to their size. However, our calculations showed an average number of long intravesicular protrusions from class 4 per SV (90 SVs per expert annotator) to be 1.0, with some vesicles having up to 4 (*SI Appendix*, Fig. S2 *A, B*). Long proteins were observed in 59% of the analyzed SVs. Due to their flexibility, we could not average them using subtomogram averaging (StA)^24,25^. The length of protrusions was between ~120-400 Å, with the most abundant value of ~180 Å. For some of them, we observed an extravesicular (cytoplasmic) part (Fig. 1 *F, SI Appendix*, Fig. S2 *A, B*).

To match the potential molecular identities for our morphologically defined protein classes 1-4, we examined AlphaFold2^22^ predictions for the proteins previously reported to be present in SVs^1,6^ (Fig. 2 *F*, *SI Appendix*, Table S2-S4). UniProt^26^ annotation of the AlphaFold2^22^ predictions, containing the information about their lumenal, transmembrane, and cytoplasmic domains, allowed us to categorize and match them with corresponding densities observed in tomograms regarding their shape and membrane topology. For example, the SV protein SV2a^27^ contains relatively large intra-and extravascular domains, resulting in the noticeable density present on both membrane sides, making it a candidate for class 2. On the contrary, Syp^28^ contains a relatively large intravesicular domain and a transmembrane domain, but no appreciable extravesicular domain, being a candidate for class 3 (Fig. 2 *F*). The set of proteins predicted as having only an extravesicular domain or extravesicular and transmembrane domains are synaptotagmins, Rab3A, synapsins, RalA, SEC22b, SCAMPs (class 1); class 2 resembles SV2 proteins, class 3 resembles Syp, synaptogyrins (Syngrs), Syp-like protein 1 (Sypl1) and synaptoporin (Synpr) (Fig.2 *F*, *SI Appendix*, Table S2-S4).

**Fig. 2.**
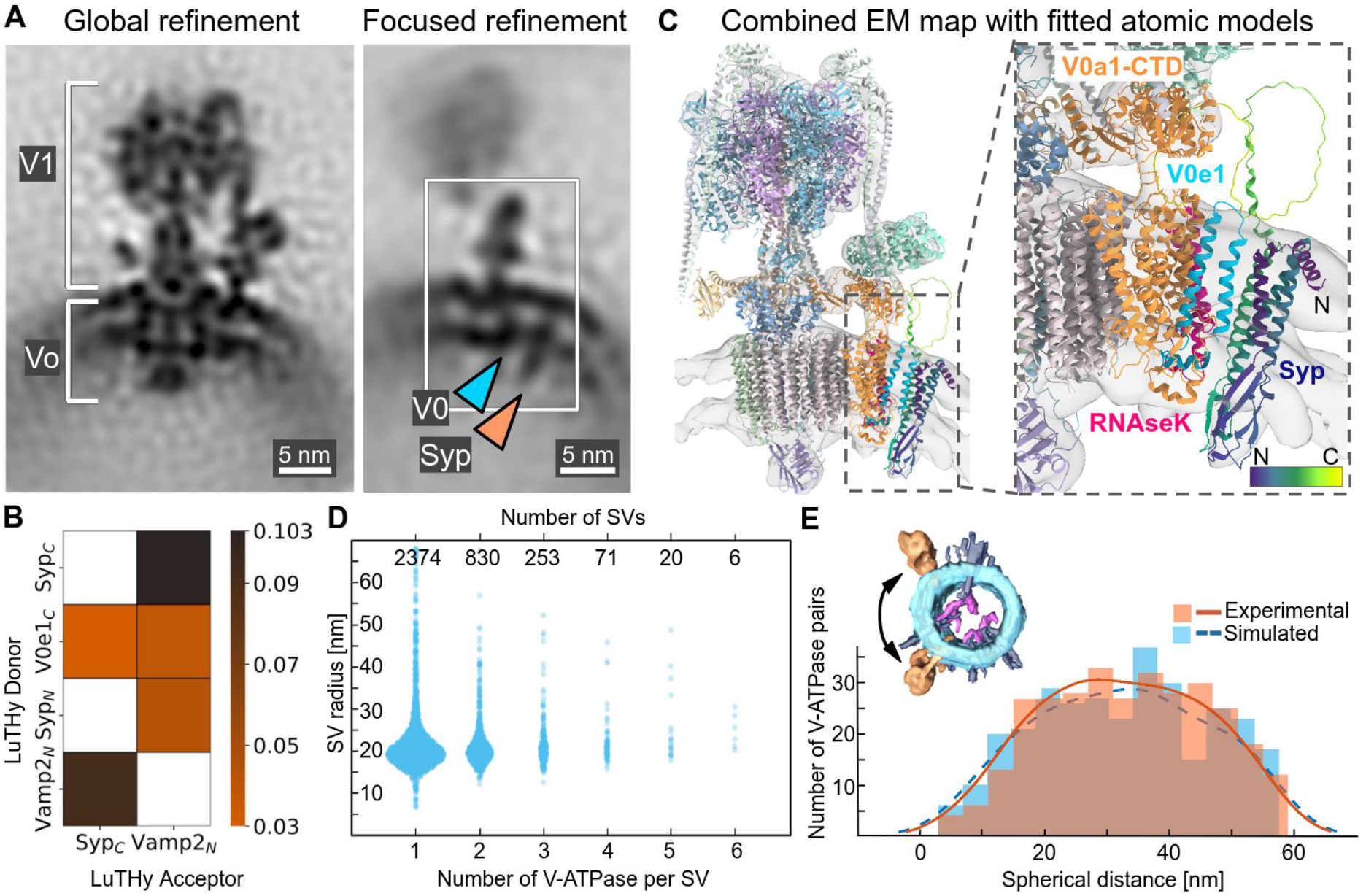
V-ATPase-Syp complex and V-ATPases arrangement on the surface of SVs. A) Left: structure of V-ATPase on the SV’s surface at 16.7 Å resolution. Right: An extra density (marked with an orange arrow) near the V-ATPase Vo region (marked with a blue arrow). Scale bars: 5 nm. B) Selected interactions (BRET signals) obtained with LuTHy assays performed with V-ATPase Voe1 domain, Syp and VAMP2. Control-corrected BRET (cBRET) ratios for the tested pairs of interacting partners as the heatmap, representing protein interaction strength with a color gradient from orange to black. Only interactions with observed cBRET values ≥0.03 (cutoff for membrane proteins) are shown. Non-tested pairs are shown by non-colored/white cells. C) Rigid-body fit of the V-ATPase and Syp atomic models into the combined map, obtained by merging global and focus-refined StA maps of V-ATPase from isolated SVs. V-ATPase Voe1 domain is shown in blue, the Voa1-CTD domain is shown in orange, RNAseK domain is shown in magenta. Syp is shown in viridis (violet-to-yellow) gradient, corresponding to the N-to-C termini. An atomic model of a purified V-ATPase complex (PDB: 6wm2) and an AlphaFold2 prediction of Syp (UniProt ID: Q62277) were used. D) Correlation of SV radii distributions, with SVs being categorized by the number of V-ATPases identified on their surfaces. E) Experimental (orange, continuous) and simulated (blue, dashed) distributions of spherical (geodesic) distances between pairs of V-ATPases on SV surfaces containing two V-ATPases. The measured distance is schematically represented with a black arrow at the volume-rendered SV.

To identify potential proteins inside the vesicles, we performed a proteinase K assay^29^. In this assay, extravesicular domains of proteins are digested by proteinase K and only proteins located within the membrane or vesicle lumen are protected. The ratio of protected proteins was quantified by label-free mass-spectrometry (*SI Appendix*, Fig. S3), showing most of the proteins to be proteinase K sensitive. Our analysis of structures predicted for SV proteins did not reveal candidates for class 4. Therefore, we examined proteins with extracellular domains that are typically located close to synaptic release sites. We analyzed the AlphaFold2 predictions for N-Cadherin (~225 Å in length)^30^, which are known to mediate cell-cell adhesion, cover an axon terminal, and undergo constant endocytosis^31,32^. Cadherins have five tandem repeating extracellular domains, a transmembrane domain, and a smaller intracellular domain^30^. Cadherin-2, 6, 10, 11, and 13 were found in our proteomics analysis (*SI*, Table S1), and cadherin 2 and 13 were protected in our proteinase K assay (*SI Appendix*, Fig. S3 *C*). Considering the size of the density and our proteomics analysis, we hypothesize that a fraction of the observed intravesicular densities could be synaptic adhesion molecules like cadherins.

### The V-ATPase forms a complex with synaptophysin

We next identified V-ATPases in our tomograms and generated a structure at a resolution of 16.7 Å from 5361 particles by StA (Fig.2 *A*, *left, SI Appendix*, Fig. S4). A previously reported atomic model of human V-ATPase from a single-particle cryo-EM analysis (PDB: 6wm2^23^) fitted well into the density. The limited number of particles without symmetry and the highly dynamic structure of the V-ATPase likely were the main resolution-limiting factors. Interestingly, in our V-ATPase structure, we observed an additional membrane-embedded density that was not accommodated by the atomic model of the V-ATPase. The density had a small SV-lumenal domain, located a few nanometers away from the Vo-a1-e-RNAseK-proximal region. A refinement focused on this area resulted in a better-defined density of ~65 Å in length with a resolution of 21.1 Å (Fig.2 *A*, *right, SI Appendix*, Fig. S4). To probe potential candidates for this density, we analyzed the proteins from class 3 (Syp, Synpr, Syngr1, Syngr3, and Syp-like protein 1) for the fit into the density, abundance per SV in the reported proteomics data^6,7^ or other interaction studies (*SI Appendix*, Fig. S5). Syp was abundant, had an overall fitting shape and was previously suggested to interact with V-ATPase^6,33^. Furthermore, Syp was found to be one of the proteinase K-protected proteins, while Syngrs and Synpr were proteinase K accessible (*SI Appendix*, Fig. S3 *C)*.

To test whether Syp interacts with subunits of the V-ATPase, we next performed a quantitative evaluation of protein-protein interactions (PPI) by the luminescence-based two-hybrid assay LuTHy^34^. LuTHy identifies direct interactions between tagged fusion proteins expressed in mammalian cells by quantification of bioluminescence resonance energy transfer (BRET) (Fig. 2 *B*, *SI Appendix*, Fig. S6, *SI* Table S2). The assay demonstrated interactions between the C-terminus of human Syp and the C-terminus of the human Voe1, an isoform of which was shown to be a subunit of the V-ATPase^23^. Furthermore, the single transmembrane helix-protein VAMP2 (synaptobrevin) interacted with both Syp and the Voe1 subunit of the V-ATPase (Fig. 2 *B*, *SI Appendix*, Fig. S6). These considerations and experiments lead to the conclusion that the additional density in the V-ATPase is accommodated by Syp, which agrees with a recent cryo-EM report^35^.

To assemble a model of the V-ATPase-Syp complex, we merged a global and a focus-refined StA maps and fitted an atomic model of the human V-ATPase from a single-particle cryo-EM analysis (PDB: 6wm2^23^) into the obtained combined map (see Methods). Further, we performed rigid-body docking of the available AlphaFold2 prediction of mouse Syp (UniProt ID: Q62277) into the same combined map (see Methods, Fig. 2 *C*). Such a model resulted in a good overall fit and did not produce clashes.

### Distribution of V-ATPases on the surface of SVs

Some V-ATPases were observed to form apparent clusters on SVs, while others seemed to be positioned away from each other (Fig.1 *E*). To quantify the distribution of V-ATPases, we manually confirmed the identity of V-ATPases using their positions and orientations obtained by StA. We matched the V-ATPases and the SVs and determined the radius of the corresponding SVs with a set of algorithms (see Methods section). 63% of SVs contained one V-ATPase/SV, 26% of SVs contained two V-ATPases/SV, 8% contained three V-ATPases/SV, and the minority contained more than three V-ATPases/SV (Fig. 2 *D*). Surprisingly, we found that the number of V-ATPases per SV does not correlate with the SV radius (Fig. 2 *D*).

To quantitatively characterize the mutual arrangement of V-ATPases on the surface of SVs, we calculated the distance between the centers of V-ATPases across the membrane, referred to as the “spherical distance” (Fig.2 *E*). For this, we only analyzed SVs within the most likely radius range of 19-20 nm with two V-ATPases present on their surfaces (n = 320 SVs). Despite the non-uniform distribution profile, its rather broad peak (~20-45 nm) appearance did not show any preferred distances between particles, while its non-uniformity is caused by the orientation bias in particle picking: V-ATPases, which are oriented in the direction of the electron beam, are underrepresented due to the “missing wedge” effect, a distortion associated with the cryo-ET imaging procedure causing an anisotropic resolution. To evaluate this, we simulated a random distribution of paired particles on the surface of SVs with the given radius, imposing the particle orientations bias, observed for our V-ATPase dataset (see Methods section). The distribution of pairwise distances reflected a non-uniform profile, similar to the experimental data, due to the accounted underrepresentation of the “top views” of V-ATPases (Fig. 2 *E*). The statistically equal distribution of measured and simulated pairwise distances (p = 0.44, two-sample Kolmogorov-Smirnov nonparametric statistical test) indicates that V-ATPases are randomly distributed on the surface of SVs without any specific clustering mechanism. We applied the same analysis to SVs with three V-ATPases and found no difference in the distributions.

### Most clathrin coats on vesicles are not fully covered

We observed fully assembled V-ATPases not only at SVs but also under clathrin cages in the *ex vivo* preparation (Fig. 3 *A*), indicating that the reassembly of the V-ATPse after SV fusion can happen early in SV recycling. The V1 heads were observed under the layer of the clathrin cage, as would be expected from the V1 region height (~19 nm) and the height between the vesicle membrane and clathrin triskelion (~24 nm, measurements in *SI Appendix*, Fig. S7 *A*). Next, we investigated clathrin-coated vesicles (CCVs) in tomograms from *in situ* neuronal terminals and in tomograms of isolated vesicles. In previous studies, clathrin symmetry variation and cage sizes were investigated in CCVs purified from brain tissue^15,16^ or assembled *in vitro*^13,14,36,37^. Here, we analyzed CCVs’ size, coverage with clathrin, and cage symmetry in purified vesicles and inside primary hippocampal neurons using cryo-ET. We found that half of CCVs in the isolated sample and a minor fraction of *in situ* CCVs contained only partial clathrin coat in the missing-wedge-free directions (Fig. 3 *B, C*, *SI Appendix*, Fig. S2 *C, D*). As the non-coated SVs, some CCVs also contained long intravesicular protrusions (Fig. 3 *B, C*, *SI Appendix*, Fig. S2 *C, D)*.

**Fig. 3.**
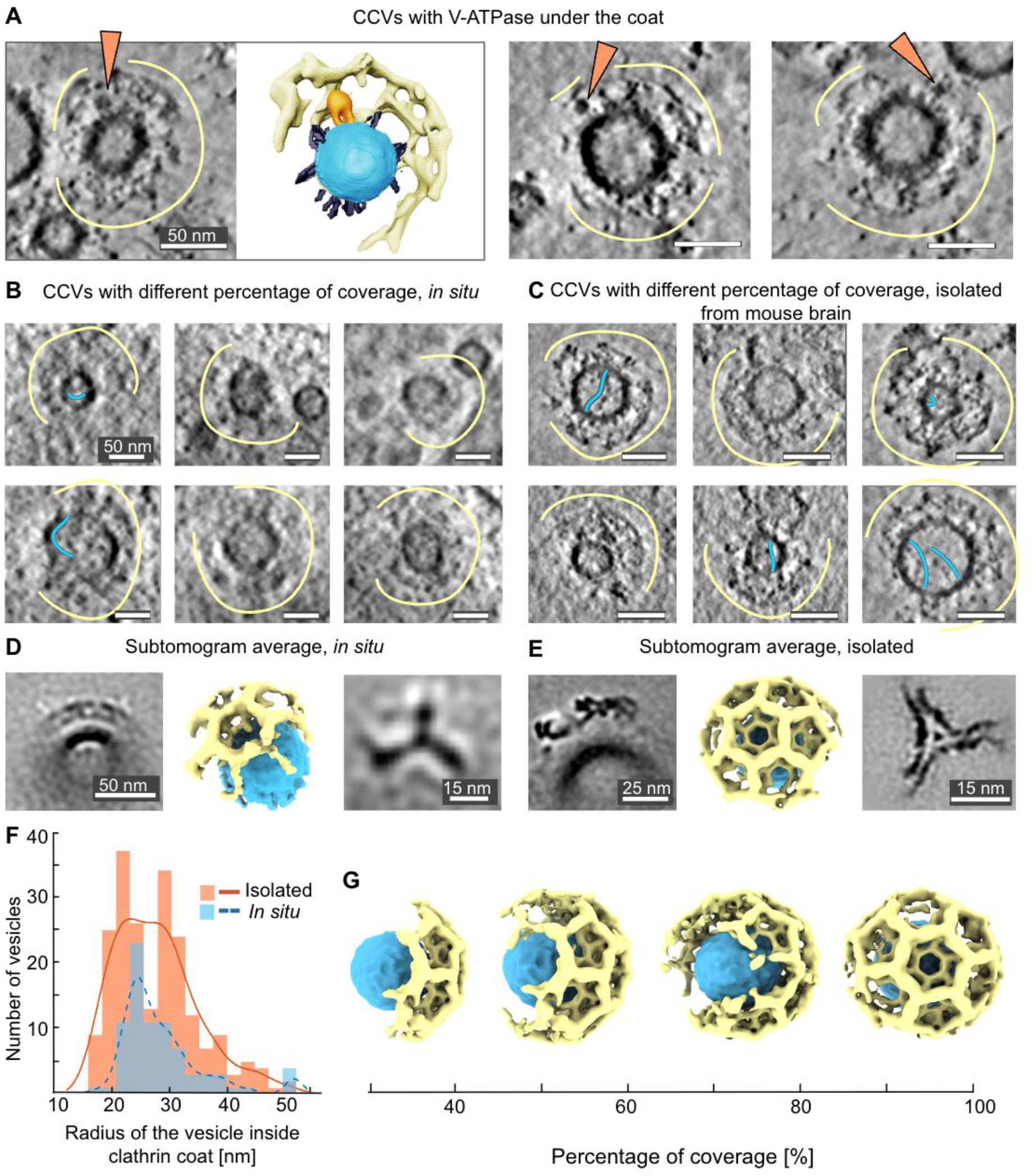
Distribution of clathrin on the surface of vesicles. A) CCVs contain V-ATPase under their cages and a segmentation of one of them. V-ATPases are shown with orange arrows, small proteins in gray, clathrin cage - with yellow curves. Segmentation: The vesicle membrane is blue, V-ATPase is orange, clathrin cage - yellow. For clarity, only half of a clathrin coat is segmented. The scale bar is 50 nm. B) CCVs from primary hippocampal neurons with different percentages of clathrin coverage. Clathrin cages are shown with yellow curves. Long intravesicular protrusions are shown with blue curves. Scale bars: 50 nm. C) CCVs, isolated from mouse brain tissue, with different percentages of clathrin coverage. Scale bars: 50 nm. D) StA of CCV fragment from primary hippocampal neurons (left, scale bar: 50 nm) with its volume rendering (ChimeraX^38^, center) and of a single clathrin triskelion (right, scale bar: 15 nm). E) StA of CCV fragments, isolated from mouse brain tissue (left, scale bar: 25 nm) with its volume rendering (ChimeraX, center) and of a clathrin triskelion (right, scale bar: 15 nm). F) Histogram of CCVs radii (CC-enclosed vesicle center-to-membrane distances) distribution, isolated from mouse brain tissue (orange, continuous) and observed in neurons (blue, dashed). G) Volume-rendered representations of individual isolated CCVs, corresponding to the different percentages of coverage with clathrin.

The knowledge of the shape and symmetry of the clathrin coat is essential for elucidating its function and regulation^15,37^. Therefore we tested the predominant arrangement of clathrin on the surface of the CCVs *in situ* and isolated vesicles. We first picked 81 CCVs from tomograms of hippocampal neurons, determined their radii (Fig. 3 *F*), parameterized their surface, and performed subtomogram classification and averaging, without the application of symmetry (C1). This resulted in a low-resolution StA structure of a clathrin coat fragment at 76 Å resolution, containing 6 triskelia (Fig.3 *D*, *SI Appendix*, Fig. S4). Most of the vesicles inside clathrin cages had radii of 25 nm (Fig. 3 *F*).

We next structurally analyzed 212 CCVs from the vesicles isolated from the mouse brain (Fig. 3 *C*). The distribution of CCV outer membrane radii showed two peaks: at 23 nm and 31 nm (Fig. 3 *F*). The observed variability of radii is likely due to the origin of the CCvs: while the vesicles from the first radii peak are possibly from synapses, the larger vesicles may be related to other cellular processes or cell types. We performed StA of the clathrin lattice containing 20 triskelia in the alignment mask, resulting in a structure at a resolution of 27 Å (Fig. 3 *E*, *SI Appendix*, Fig. S4). The structure showed a hexamer surrounded by 3 pentamers and 3 hexamers (soccer ball) cage symmetry (Fig. 3 *E*). Thus, in both isolated CCVs and vesicles *in situ*, we observed a soccer ball symmetry as the most dominant type of clathrin arrangement, independent of the vesicle radii.

### Synapses contain clathrin baskets without vesicles inside

We observed clathrin baskets without vesicles inside both *in situ* and in the isolated vesicle preparations (Fig. 4 *A*, *SI Appendix*, Fig. S8). In neurons, empty cages were observed at postsynapses and nonsynaptic neurite compartments (Fig. 4 *A*). However, clathrin baskets were preferentially located in the proximity of membranes (shortest distance to membrane: modal value 70 nm, median 130 nm; Fig. 4 *B, C*, *SI Appendix*, Fig. S8 *C*). Calculated individual cage-to-membrane distances differed significantly from a simulated dataset mimicking a uniform distribution of empty cages within cellular compartments (p = 0.007, Kolmogorov-Smirnov test), suggesting that the empty baskets tend to be positioned closer to the membranes than would be expected randomly (see Methods).

**Fig. 4.**
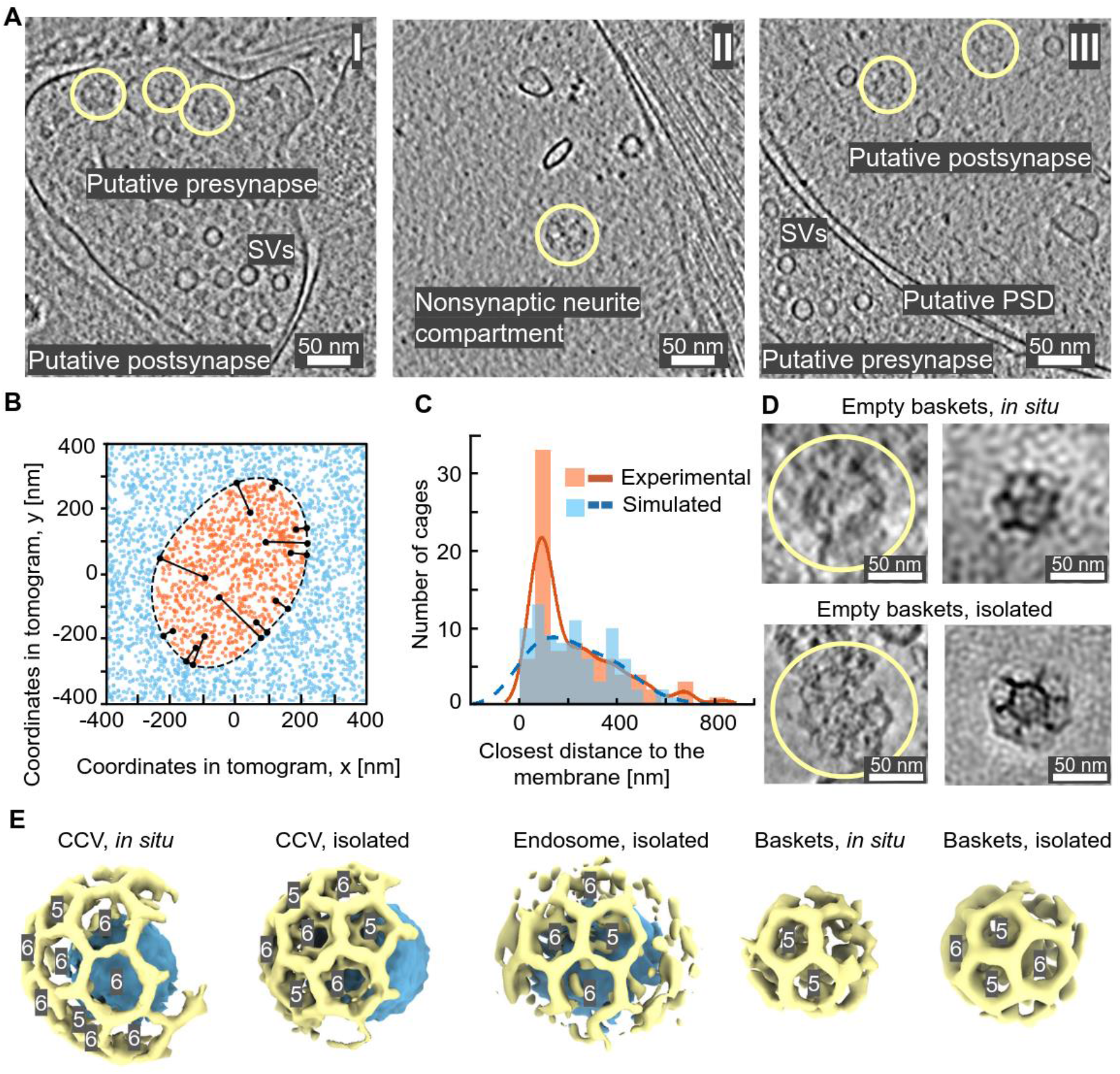
Non-vesicle-containing clathrin baskets are present in cells. A) Slices through tomograms with empty clathrin baskets from the *in situ* samples indicate their localization in presynapse (I), nonsynaptic neurite compartment (II), and postsynapse (III). Clathrin baskets are marked as yellow circles. B) Monte Carlo simulations of clathrin cages uniformly distributed in cells. The modeled synapse is indicated with a dashed line, orange-intracellular, blue - extracellular space. Black lines inside indicate the distances between the simulated randomly distributed dots (representing cages) and the membrane. C) Clathrin-basket-to-membrane distances, experimental (orange, solid) and simulated (blue, dashed) distributions. D) Top: Clathrin baskets as seen in cryo-ET volumes from on-grid-grown neurons and their StA. Clathrin cages are shown with yellow curves. Bottom: isolated clathrin baskets and their StA. E) Clathrin cages are shown to scale for isolated and *in situ* CCVs, endosomes, and empty clathrin baskets. Scale bars A, D, E: 50 nm.

The average radius of 149 empty clathrin cages from mouse brain tissue was ~36 nm, which is significantly smaller than clathrin cages with a vesicle inside (Fig. 4 *D, E*, *SI Appendix*, Fig. S8 *B*). Ninety-two empty clathrin cages from on-grid-grown neurons had radii of 30-38 nm. We performed StA for empty clathrin cages in both datasets (*in situ* and isolated vesicles), and determined low-resolution structures (Fig. 4 *D*, *SI Appendix*, Fig. S4), showing disordered density inside the cages (*SI Appendix*, Fig. S8 *D*). The structure obtained from the brain preparation contained a hexamer interacting with four pentamers and one hexamer, and one unclearly-resolved polygon, potentially C2-basket geometry^15^ (marked in *SI Appendix*, Fig. S8 *D*).

Furthermore, we observed large ellipsoidal vesicles partially coated with clathrin, resembling endosomes, both *in situ* and in the isolated vesicle preparation (*SI Appendix*, Fig. S7 *B*). We performed StA of the membrane segments containing clathrin coats using the available number of particles (n = 27, *SI Appendix*, Fig. S4). The resulting structure had a soccer-ball symmetry with 1 pentamer, 4 hexamers and 1 unclear polygon (Fig. 4 *E*, *SI Appendix*, Fig. S4). Thus, our structures of empty clathrin cages showed a different symmetry type compared to the clathrin coats containing a vesicle inside or located at endosomes.

## Discussion

In this study, we combined *in situ* and *ex vivo* cryo-ET with biophysical methods, and structural and statistical analysis to perform a structural and morphometric characterization of proteins associated with SVs and CCVs. SVs isolated from mouse brains and SVs within neurons cultured on EM gids showed a comparable overall morphology, mass spectrometry confirmed a high abundance of SV proteins in our isolated vesicle preparation. Due to the lower sample thickness, our *ex vivo* SV dataset provided a higher signal, which allowed us to characterize and classify proteins integrated into the SV membrane and located inside the SV lumen. Structural analysis of these proteins involving StA has not been possible because of their heterogeneity and small size: VAMP2 - 18 kDa^39^, Syp - 38 kDa^40^, synaptotagmin-1 - 65 kDa^41^. A larger size would be required for structural analysis: recently, a structure of a 120 kDa^42^ mostly membrane-embedded protein was reported at an intermediate resolution of ~16 Å^42^. The highly abundant SV proteins such as Syp and neurotransmitter transporters are membrane-embedded, which further limits their detection by cryo-ET, creating the observed apparent empty membrane. Nevertheless, we were able to classify observed protein densities into four categories and matched them with AlphaFold2 predictions of known SV proteins. Beyond that, we identified long intravesicular protein densities. Since none of the SV proteins previously identified via mass spectrometry or fluorescence microscopy^1,2,4,6^ fitted into these densities, we suggest that extracellular adhesion proteins like cadherins account, at least in parts, for these densities. A possible explanation for their intravesicular localization is that adhesion molecules are internalized during endocytosis, which is supported by our finding of long protrusions also inside CCVs. SVs may transport adhesion molecules toward the synapse, where they have been observed^43^.

Even though intact SVs display a convoluted matrix with high protein density, we were able to characterize the abundance, distribution, and structure of V-ATPases. We showed that SVs had an average of 1.4 V-ATPases per SV, randomly distributed on their surface, suggesting that the positioning of V-ATPases is not crucial for their vesicular function. We produced an intermediate-resolution structure of the V-ATPase by StA and observed an ordered density next to its transmembrane domain, potentially forming a functional complex. We attribute the observed density containing a transmembrane segment and a small intravesicular domain to the protein Syp for the following reasons: (i) fitting of several candidate proteins to the density yielded best results for Syp, synaptoporin and synaptophysin-like protein 1, (ii) our proteinase K assay indicated that Syp is highly protected, supporting its mostly membrane-embedded and lumenal localization (*SI Appendix*, Fig. S3 *C*), (iii) our LuTHy assay showed an interaction between human Syp and the V-ATPase Voe1 subunit. Independent from our study, a recent high-resolution structure of the rat V-ATPase suggested an interaction with Syp^35^. Furthermore, previous cross-linking MS data indicated direct interactions between Syp and V-ATPase and between VAMP2 and both Voa1 and Syp. Thus, it is likely that VAMP2, Syp, and the V-ATPase Vo region form complexes, as already suggested before^33^. Although it was not possible to position VAMP2 in our structure due to its small size of 18 kDa and a potentially less stable interaction, our LuTHy assay corroborated the presence of such a triumvirate, as it demonstrated strong pairwise interactions between VAMP2 and Voe1 and Syp.

The interactions between V-ATPase, Syp, and VAMP2 may constitute a control mechanism to generate and maintain fully functional SVs over the rounds of SV recycling^44^. Mice lacking the SNARE protein VAMP2 are severely burdened because synaptic exocytosis is almost completely blocked^45^. Likewise, acute photo-inactivation of V-ATPase subunits was shown to strongly impair neurotransmitter release^46^. In contrast, synaptophysin knockout mice have a normal life expectancy and only show mild behavioral alterations^40,47^. Syp-knockout mice, as well as humans harboring mutations in the synaptophysin gene, display learning deficits and intellectual disabilities^47,48^. While Syp is not required for baseline neurotransmission^49^, a stronger short-term depression can be observed during sustained synaptic transmission in the absence of Syp^50^. This observation has been explained by the impaired Syp-dependent retrieval of VAMP2 during endocytosis and SV reformation^49,50^. Taking into account that VAMP2 and Syp are the two most abundant proteins on SVs, whereas only 1.4 copies of the V-ATPase are present per SV on average, but that all three proteins interact with each other, it may be questioned if the observed phenotype in absence of Syp is indeed only caused by the decreasing prevalence of VAMP2. Instead, Syp may function as a scaffold securing not only the proper recycling of VAMP2 but also of at least one V-ATPase per SV. In this scenario, sustained exocytosis in the absence of Syp would lead to an accumulation of nonfunctional SVs either lacking V-ATPses and thus neurotransmitters or VAMP2.

While the Vo region of the V-ATPase remains membrane-embedded during exocytosis and SV recycling, the V1 region is supposed to be disassembled short before fusion and needs to be reassembled. Previous studies indicated that V1 region proteins might be detected at the plasma membrane and at CCVs^18,20^. With our cryo-ET analysis of *ex vivo* CCVs, we now confirmed that the Vo and V1 may assemble at CCVs, indicating that the V1 region may be recruited already at an early stage of SV reformation. Our investigation of CCVs further revealed that not all clathrin coats were fully assembled. We identified partially uncoated CCVs both *in situ* and in brain preparations. Notably, clathrin coats in most of the observed CCVs had a soccer-ball symmetry. We propose that these vesicles initially arise from the fission of clathrin-coated pits but subsequently undergo stepwise uncoating, which typically involves the proteins synaptojanin, Hsc70, and auxilin and is ATP-dependent^51,52^. Importantly, we observed most CCVs i*n situ* in extrasynaptic parts of neurites. The lower abundance of CCVs at the presynapse is likely a consequence of the fast SV turnover, with exo- and endocytic proteins being enriched close to the synaptic release sites. In contrast, activity-independent CME taking place at endocytic zones farther away from the presynapse is supposed to be slower and only promoted when necessary^53^. Thus, we were able to capture the intermediate stages of the uncoating process enriched close to these endocytic zones. Since we did not detect any differences in the size, organization, or symmetry of CCVs from presynapses versus post-or extrasynaptic regions, we expect that the individual steps of CME are conserved between different neuronal compartments and different cellular functions.

The role of the observed empty clathrin baskets in synapses remains unclear. Studies have demonstrated that clathrin can be disassembled and reassembled into cages of different symmetry *in vitro*^54^ and in cells^55,56^. The empty baskets may thus arise from clathrin polymers pre-assembled without a vesicle inside. These preassembled baskets could display a clathrin reservoir close to endocytic zones to facilitate the fast recruitment of clathrin triskelia or even partial lattices. Alternatively, the empty clathrin baskets may result from incomplete vesicle uncoating: since the vesicle uncoating initiates at hexagon corners^15^, the relative abundance of pentamers increases over time. This may facilitate the formation of C2-symmetry baskets. Whether these baskets display a reservoir for faster clathrin recruitment and assembly or whether they are deposited for subsequent degradation, remains to be examined further.

In conclusion, our analysis provided new insights into the molecular architecture of synaptic vesicles and the mechanisms of their assembly and recycling. Further progress in understanding the molecular landscapes of individual SVs may be expected from the ongoing developments in cryo-ET hardware, such as phase plates and improvements in detectors and software. Furthermore, combinations of cryo-ET with labeling techniques and complementary information about protein-protein interactions may provide further insights into the molecular architecture of SVs, their functional parts, and associated diseases.

### Animals

All experimental procedures involving the use of mice were carried out in accordance with national and institutional guidelines. C57/BL6N wildtype mice for SV preparations in the study were bred at the Leibniz Research Institute for Molecular Pharmacology. Hippocampal primary culture neurons were prepared from P0-P2 C57/BL6N wildtype mice provided by Charité-Universitätsmedizin Berlin.

## Methods

### Culture of primary hippocampal neurons

Preparations of astrocytes and neurons from P0-P2 mice were performed as described previously^57^. A feeder layer of astrocytes isolated from mouse cortices of either sex was seeded on collagen/ poly-D-lysine coated 6-well plates and cultured for 1-2 weeks at 37°C in DMEM medium containing 10% fetal calf serum (FCS) and 1% penicillin/streptomycin. Glia proliferation was arrested by the addition of 5-fluoro-2-deoxyuridine (8 μM) and uridine (20 μM) overnight and the medium was subsequently replaced by Neurobasal-A medium supplemented with B-27, Glutamax and penicillin/streptomycin prior neuron seeding. Quantifoil R3.5/1 AU holey carbon 400 mesh grids were glow-discharged, coated with collagen/poly-D-lysine, and placed on pedestals on top of 1-2 weeks old astrocytes. Primary hippocampal neurons were plated with a density of 150,00-200,000 cells/well and grown for 17 days at 37°C until further use.

### Synaptic vesicle enrichment from mouse brain tissue

Vesicles from murine neurons were isolated as described previously^1,58,59^, stopping after LP2 fraction. All steps were performed at 4 °C. Briefly, the hippocampus, cortex, and cerebellum of six 40-days-old mice were extracted and homogenized in homogenization buffer (HB; 4 mM HEPES pH 7.4, 320 mM sucrose, 1 mM PMSF, protease inhibitor mix) with 9 strokes in a Dounce homogenizer at 900 rpm. Tissue debris was removed by centrifugation at 800xg for 10 min. The supernatant was centrifuged at 10,000xg for 15 min and the pellet containing synaptosomes was collected and washed with HB. Synaptosomes were swelled in a hypotonic buffer (1:10 HB in H_2_O) and lysed with three strokes in the Dounce homogenizer at 2,000 rpm. Isotonic conditions were restored and the mixture was centrifuged at 25,000xg for 20 min. Subsequently, the supernatant was centrifuged at 200,000xg for 2 h. The pellet was resuspended in 150 μL HB and the protein concentration was determined by Bradford assay and negative staining.

### Negative staining of murine synaptic vesicle preparations

Synaptic vesicle preparations were diluted 1:100 (v/v) in PBS and 4 μL of the solution was applied to a glow-discharged continuous carbon grid. The sample was incubated on the grid for 30 s and washed with water for 30 s. Then, the grid was incubated with 2% uranyl-acetate for 30 s. The grids were imaged on a Zeiss 910, FEI Morgagni.

### Proteinase K treatment of synaptic vesicles and proteolytic digestion

The PK-sensitivity assay was carried out as previously described^29^. In brief, synaptic vesicles were diluted in HB (without protease inhibitors) supplemented with 5 mM CaCl_2_. In three replicates, vesicles were incubated with or without 50 μg/mL Proteinase K (PK) for 1 h at 37 °C. PK was inhibited with 5 mM PMSF for 10 min at RT. Proteins and polypeptides were reduced, alkylated, and digested using Lys-C (1:200 w/w) and trypsin (1:100 w/w). The digestion was stopped with 1% FA and peptides desalted with C18-StageTips. Proteomic samples for the determination of protein copy numbers and sample purity and composition were processed similarly but without the use of PK.

#### LC-MS/MS analysis

Separation of the samples was achieved by reverse phase (RP)-HPLC on a Thermo Scientific™ Dionex™ UltiMate™ 3000 system connected to a PepMap C-18 trap column (Thermo Scientific) and an in-house packed C18 column for reversed-phase separation at 300 nL/min flow rate over120 or 180 min. Samples were analyzed on an Orbitrap Fusion or Orbitrap Fusion Lumos mass spectrometer with Instrument Control Software version 3.4. Data was acquired in data-dependent acquisition (DDA) mode. MS1 scans were acquired in the Orbitrap with a mass resolution of 120,000. MS1 scan range was set to m/z 375 – 1,500, 100% normalized AGC target, 50 ms maximum injection time, and 40 s dynamic exclusion. MS2 scans were acquired in the Ion trap in rapid mode. The normalized AGC target was set to 100%, 35 ms injection time, and an isolation window 1.6 m/z. Only precursors at charged states +2 to +4 were subjected to MS2. Peptides were fragmented using Normalized Collision Energy (NCE) of 30% or 32%.

#### Mass spectrometry data analysis

RAW files were searched using MaxQuant^60^ (Version 2.0.3.0) using the following parameters: MS1 mass tolerance, 10 ppm; MS2 mass tolerance, 0.5 Da; the maximum number of missed cleavages, 2; minimum peptide length, 7; peptide-mass, 500 – 5,000 Da. Carbamidomethylation (+ 57.021 Da) on cysteines was used as a static modification. Oxidation of methionine (+15.995 Da) and acetylation of the protein N-terminus (+42.011 Da) were set as a variable modification. Data were searched against the murine proteome retrieved from UniProt^26^. Label-free quantification^60^ and iBAQ calculation were performed with a match between runs activated. Volcano plots were generated using the EnhancedVolcano v1.14.0 R package. Subcellular locations were retrieved from the Uniprot and transmembrane domains predicted using deepTMHMM^61^.

#### Sample preparation for cryo-electron tomography

Four biologically replicated sample preparations, 20 μl of the vesicles sample (c = 0.15 mg/ml) were mixed with five-or ten-nanometer gold fiducial markers (purchased from the Cell Microscopy Core, Utrecht University) with the ratio of 1:10. Four μl of this mixture was deposited to glow-discharged cryo-EM grids (UltrAuFoil R1.2/1.3, Quantifoil Cu 2/2, or Quantifoil AU 100 Holey carbon Films R2/1), blotted for 3 seconds at relative humidity 98% and temperature 4°C and plunge-frozen into liquid ethane (Vitrobot). Three types of samples were prepared - one without additional factors, the second one with V-ATPase inhibitor bafilomycin A1 to prevent proton translocation through V-ATPase, and the third one - with ATP. The ATP-containing cryo-sample was prepared as described before^62^. Briefly, 3 μl of freshly isolated SVs were mixed with 1 μl of ATP solution in PBS containing MgCl2 (final concentration of ATP c = 4.5 μM, final concentration of MgCl_2_ c = 5 μM), and incubated on ice for 5 minutes before applying on the grid. The bafilomycin A1-containing cryo-sample was prepared as described previously^63^. A separate aliquot of SV was incubated with bafilomycin A1 (final c = 90 nM) on ice for 15 minutes prior to freezing. The initial approach was to perform subtomogram classification to obtain several conformational states of V-ATPase in the native environment. However, the achieved resolution for each of the three datasets (?25 Å) did not allow us to detect differences. Therefore, all datasets were processed together, aiming at separating V-ATPase particles from false positive particles, followed by the analysis of V-ATPase distribution over vesicles and potential clustering. For the neurons grown directly on grids, a freezing solution was prepared containing 140 mM NaCl, 2.4 mM KCl, 10 mM HEPES, 10 mM glucose, 4 mM CaCl_2_, 1 mM MgCl_2_, 3 μM NBQX, and 30 μM bicuculline (~300mOsm; pH7.4). 10 nm BSA-gold in freezing solution (OD~2) was used as fiducial markers. Grids were briefly washed in freezing solution pre-warmed to 37°C, then transferred to the plunge freezer (Vitrobot). 4 μl fiducial marker-containing freezing solution was added to the grid prior to blotting for 16 s (backside blotting, blot force 10) at 37°C and 80% relative humidity and plunge-freezing into liquid ethane.

#### Cryo-ET data collection, image processing, and subtomogram averaging

Three dose-symmetrical and four dose-asymmetrical^64^ data collection sessions were performed on a TFS Titan Krios electron microscope operated at 300 kV with a Gatan Quantum energy filter with a slit width of 20 eV and a K3 (Gatan) direct detector operated in counting mode. For several data collection, different total exposures were used: ~128 e^−^/Å^2^, ~270 e^−^/Å^2^, ~229 e^−^/Å^2^, ~271 e^−^/Å^2^, ~213 e^−^/Å^2^ was equally (for 3 datasets) ^65^, or unequally (for 4 datasets)^66^ distributed between 31, 35 or 37 tilts. Ten frame movies were acquired for each tilt. The details of data collection are given in Table S2. The number of collected tomograms: hippocampal neurons - 88 tilt series were collected (superresolution pixel size = 3.07 Å) using PACE-tomo^67^; for isolated SV fractions - 719 tilt series (superresolution pixel size = 0.84 Å), 438 of which were collected with PACE-tomo^67^.

Data processing was streamlined using TomoBEAR^68^. The aligned frames were motion-corrected using MotionCor2^69^. Tilt-series alignment was performed by DynamoTSA^70^ and manually inspected and refined in IMOD^71,72^, using the 10-nm or 5-nm gold fiducial markers. For each projection, the defocus values were measured by Gctf^73^, and CTF correction was performed using ctfphaseflip^74^ from IMOD. Weighted back-projection in IMOD was used to produce sixteen times binned reconstructions of 719 tomograms generated from CTF-corrected, aligned stacks. Particles were picked manually (V-type ATPases and clathrin baskets) from 16-times binned super-resolution tomograms collected from the brain tissue (voxel size: 13.44 Å), with further extraction of 60×60×60 sized boxes (V-ATPase) and 116×116×116 sized boxes (empty clathrin baskets).

For V-ATPase, three types of datasets were recorded - one without additional factors (450 tomograms), one with an inhibitor bafilomycin A (179 tomograms), and one with additional ATP (90 tomograms).

For structural analysis, 500 particles were used to perform the initial manual alignment in *dynamo_gallery*. Roughly aligned particles were averaged to produce an initial average which was used as an initial reference to perform the global alignment of all subtomograms. Several rounds of multireference alignment and classification were performed in Dynamo, leading to the removal of suboptimal particles. Classification rounds were performed in several iterations, with the first range of 360° around the unit sphere. C1 symmetry was used and a wide soft-edged cylindrical mask. Aligned particles in good and bad classes were examined manually in *dynamo_gallery* to minimize the loss of good particles, which was important for further statistical analysis. In total, 5361 particles were identified as “good” ones and transferred to RELION-4.0^75^ for further refinement and 3D classification, using 4-times binned super-resolution stacks and a protein-shaped mask created in ChimeraX^38^. The final 3D average originating from 5361 particles was refined to a resolution of 16.7 Å resolution. During processing, no symmetry was applied. We did not observe differences between the structures reconstructed from different datasets; the highest resolution structure was obtained by combining all three datasets. The same final particle set was used for further focused refinement on the Vo-a1 domain with the soft-edged cylindrical mask. This resulted in the 21.1 Å resolution map with a membrane protein density with a short intravesicular domain.

For empty clathrin baskets, both in cells and isolated ones, a similar Dynamo pipeline was used. The particles were picked manually. Sixteen-times binned super-resolution subtomograms were refined and 3D-classified using a spherical soft-edged mask in Dynamo. This resulted in low-resolution structures: 90Å for the cage isolated from mouse brain tissue (51 baskets), and 160 Å for the cage from primary hippocampal neurons (34 baskets).

Sixteen-times binned super-resolution CCVs’ subtomograms were picked using a spherical model in the Dynamo Catalogue system^76^, with the cropping mesh of 5 voxels and further extraction of 64×64×64 sized boxes. Each subtomogram contained a piece of membrane and a fragment of a clathrin cage. Further, 3D-refinement and Dynamo classification resulted in 1702 particles. The same pipeline was applied to isolated endosomes, resulting in 27 particles.

For clathrin cages, obtained from primary hippocampal neurons, the same pipeline was applied. Due to the higher voxel size, the cropping mesh of 3 voxels was used, and further extraction of 60×60×60-sized boxes. Finally, 1564 particles were refined in Dynamo with a final resolution of 75.6 Å.

To perform a single triskelia refinement both for CCVs from brain tissue and from primary hippocampal neurons, the following pipeline was applied: structures, obtained from the final CCVs classification round, were centered on the clathrin hexamer, with C6 symmetry applied. That made it possible to crop single triskelia. Further refinement of single-triskelion was performed in Dynamo. The final amount of particles for cellular data was 3658 with a resolution of 81.1 Å (voxel size of 24.56 Å). The final number of triskelia particles that originated from brain-isolated CCV was 8006 (voxel size of 13.44 Å). Particles, containing triskelia from isolated mouse brain tissue, were imported in RELION-4.0^75^ and further refined and classified, which led to the 15.6 Å resolution structure (voxel size of 3.36 Å).

#### Visualization and rendering of cryo-ET volumes

Volume rendering of subtomogram averages of V-ATPAse and CCV (main text), as well as clathrin coats in the Supplementary Figures, was prepared using ChimeraX^77^. Volume segmentation and rendering of SV surfaces with embedded V-ATPases (Fig. 1 *E*), CCV with a V-ATPase under the cage (Fig 3 *A*), and clathrin-coated endosomes (*SI Appendix, Fig. S7*) were prepared using AMIRA (Thermo Fisher Scientific and Zuse Institute Berlin). The raw volume slices of tomograms, subtomograms, and subtomogram averages were obtained using IMOD^71^, Dynamo^70^, or RELION-4^75^. Schematic representation of used sample preparation procedures in Fig. 1 *A, C* are made with Biorender.

#### Protein-protein interaction assay by LuTHy

Targeted validation of Syp interactions with LuTHy assays was performed as described in Trepte et al^34^. Briefly, open reading frames of candidate human SV proteins interactors (Syp, V-ATPase Voe1, V-ATPase Voa1, VAMP2) were cloned into LuTHy expression vectors (*SI* Table S2) by standard linear recombination reactions using the Gateway Cloning System and validated by restriction enzyme digests, agarose gel electrophoresis, and Sanger sequencing. All possible eight orientations were tested for interaction pairs: N/C-terminal tagging with Nano-luciferase (NL) or Protein A-mCitrine (PA-mCit). LuTHy control vectors expressing only NL or PA-mCit were used for the calculation of corrected scores. HEK293 cells were reverse transfected using linear polyethyleneimine (25 kDa, Polysciences 23966) and LuTHy constructs; cells were subsequently incubated for 48 h. In-cell BRET measurements were carried out in flat-bottom white 96-well plates (Greiner, 655983) with 24 PPIs per plate (each PPI in triplicate). For cell-free LuC measurements, cells in 96-well plates were lysed and lysates were transferred to 384-well plates resulting in 96 PPIs per plate (Greiner, 784074). InfiniteÒ microplate readerM1000 (Tecan) was used for the readouts with the following settings: fluorescence of mCitrine recorded at Ex 500 nm/Em 530 nm, luminescence measured using blue (370–480 nm) and green (520–570 nm) bandpass filters with 1,000 ms (LuTHy-BRET) or 200 ms (LuTHy-LuC) integration time. A PPI was considered positive if its corrected BRET (cBRET) ratio was ≥ 0.03.

#### Characterization of synaptic vesicles and arrangement of V-ATPases

The identified SVs were characterized by their radii and the number of identified V-ATPases, observable (picked) on their surface. The described analysis was performed in MATLAB.

To estimate SV radii, the SV-containing subtomograms were cross-correlated with a set of binary vesicle-like masks of different radii. Each mask consisted of two encapsulated concentric spherical shells, imitating contrast produced by membrane leaflets. The cross-correlation allowed us to estimate the radii of SVs and to refine the coordinates of their centers.

Next, to calculate the number of V-ATPases and the pairwise distances between them, the subtomograms of the picked V-ATPases were first matched with the corresponding subtomograms of the SVs. The assignment was made based on the corresponding geometrical constraints using refined coordinates of vesicle centers and picked V-ATPases as well as ATPases orientations. For that, for each pair of SV and V-ATPase subtomogram, two vectors were constructed and compared:

- SV/V-ATPase vector - a vector from the refined vesicle center to the picked (presumably) V-ATPase position;
- V-ATPase vector - a unit vector representing the orientation of the V-ATPase in the membrane.

The SV/V-ATPase vector length was compared to the previously identified SV radius in order to reject picked particles that do not belong to a certain SV. Using the maximally allowed error of 5 voxels (voxel size = 13,44 Å), the V-ATPase particles were pre-assigned to the corresponding synaptic vesicles. Further, SV/V-ATPase vector orientation was compared to the pre-assigned V-ATPase vector. If the absolute angular difference between the directions of those two vectors was not exceeding 5 degrees, the V-ATPase was finally associated with the current synaptic vesicle. Using the V-ATPases/SVs matching information, the correspondence between the number of observable V-ATPases per SV and the measured SV radius was further analyzed.

#### Simulation of randomly distributed V-ATPase pairs on the SV surface

To assess whether the observed distribution of the pairwise spherical distances between V-ATPases reflects their random allocation on the SV surface, the corresponding simulated dataset was generated. During those simulations, we took into account the orientation bias associated with the electron beam direction relative to particle orientations. The described simulation was performed in MATLAB.

The further described simulations utilized the following coordinate systems: Cartesian, spherical, and cylindrical. As a main XYZ frame, we decided to use a left-handed Cartesian coordinate system. For the other coordinate systems, the following conventions have been used:

a. spherical: the radial distance r_sph_∈[0;+∞], azimuthal angle θ∈[0;2π] counted from axis X, and polar angle ϕ∈[0;π] counted from axis Z of the XYZ frame;
b. cylindrical: radial distance r_cyl_∈[0;+∞], azimuthal angle θ∈[0;2π] counted from axis X of the XYZ frame, and height z∈[-∞;+∞].

First, to imitate a set of randomly allocated V-ATPase pairs on the SV surface, the points have to be sampled uniformly on the unit sphere surface. Note that independently sampling azimuthal angle θ from uniform distribution U(0,π) and polar angle ϕ from U(0,2π) would lead to the more dense arrangement of simulated points on the poles in comparison with the equator, making the resulting distribution non-uniform. In order to generate a uniform (equidistant) distribution of points on the sphere surface, we used the possibility of establishing a geometrical mapping between the sphere surface and the cylinder side surface (i.e. excluding its caps) circumscribed around that sphere. Thus, the target uniform distribution of the sphere surface points is achieved by generating their cylindrical coordinates - azimuthal angle θ from U(0,π) and height z from U(−1,1) - and mapping them back to the sphere surface.

Further, to consider the observed orientation bias of particle picking in the simulation, the corresponding orientation angles of the generated particles have to be sampled from the respective empirical distribution. To achieve that the following steps were performed:

1. The empirical distribution of spherical polar angles ϕ_emp_∈[0;π], representing orientation angles of particles relative to the beam direction, was mapped to the respective cylindrical height coordinates z_emp_∈[-1;1] using z_emp_ = cos(ϕ_emp_).
2. The corresponding empirical cumulative density function (eCDF) for the cylindrical height values z_emp_ distribution was generated, inverted (eCDF^-1^) and equidistantly interpolated with the step of 0.01.

The obtained interpolation table of eCDF^-1^ provides a mapping of uniformly generated numbers U(0,1) to the range of simulated cylindrical heights z_sim_∈[-1;1], allowing to account for the orientation bias while generating particle coordinates.

Finally, the target V-ATPases distribution was generated following the next procedure:

1. The simulated azimuthal angles θ_sim_∈[0;2π] were sampled from the uniform distribution U(0,2π).
2. The simulated cylindrical height values z_sim_∈[-1;1] were generated by mapping points sampled from U(0,1) using the obtained interpolation table of eCDF^-1^.
3. To obtain the target simulated particle coordinates set, the generated cylindrical coordinates are transformed back to the spherical coordinates.

The described above approach allowed us to simulate the set of coordinates of V-ATPase pairs randomly allocated on the SV surface, taking into account the orientation bias. Based on the obtained pairs of particle coordinates, simulated on the unit sphere, the corresponding spherical (geodesic) distances between the V-ATPases were calculated and scaled according to the most abundant SV radius values of 19-20 nm. The number of simulated particle pairs was set to n=320, which is the number of experimentally observed vesicles containing only two ATPases within the described SV radius range.

#### Synaptophysin/V-ATPase complex model assembly

To assemble a model of the V-ATPase-Syp complex, we first combined the obtained global and focus-refined EM maps into a single one using available UCSF ChimeraX tools: *fitmap* - to superimpose initial global and focused maps, *vop maximum* - to merge them into a single ‘fused’ map. Further, we used the previously deposited purified human V-ATPase complex (PDB: 6wm2) and rigid-body fitted it (ChimeraX) to the obtained ‘fused’ V-ATPase StA map from mouse brain tissue. Next, using the Match-Maker tool in ChimeraX we have fitted the available AlphaFold2-predicted atomic model of mouse Syp (UniProt ID: Q62277) into the ‘fused’ StA map near the Voa1/Voe1/RNAseK subunits of the previously fitted V-ATPase. This allowed us to build the full initial atomic model of the Syp-V-ATPase complex (Fig. S5).

#### Simulation of randomly distributed empty clathrin cages inside cells

To assess whether the observed distribution of the distances from empty clathrin cages to the closest synaptic membrane surfaces reflects their random allocation in the neuronal cells, the corresponding simulated dataset was generated.

To account for a variety of observed neuronal cell morphologies, containing empty clathrin baskets, we created 15 surface models by manual annotation of the cellular membranes using Dynamo Catalogue tools^76^. The obtained surface models were projected onto a 2D plane and the corresponding point clouds were thinned by calculating the moving-average positions of the sub-clouds of 10 dots sorted according to their polar angles. The obtained single-dot-width point curves were fitted using B-splines of the third order. The resulting set of enclosed curves served as the cell membrane models, reflecting the distribution of cell sizes and shapes.

Further, for each cell model, we have generated 5000 randomly distributed particles, representing empty clathrin cages, sampling their coordinates from a 2D uniform distribution. Using the Monte Carlo approach and computational geometry tools, we have identified the particles located inside cell models and calculated their distances to the closest cell model surfaces. That yielded the target random distribution of the simulated empty clathrin cages inside cells.

Finally, the simulated and experimental distributions of the distances from empty clathrin baskets to the closest synaptic membranes were compared using a two-sample Kolmogorov-Smirnov nonparametric test. The probed null hypothesis was the cumulative distribution function of the experimental data being greater than for the simulated data. The obtained *p*-value of 0.007 < 0.05, suggests that the null hypothesis should be rejected in favor of the experimental data distribution being shifted towards lower values in comparison with the simulated data.

All the described calculations in this section were made using Python using open-source packages for data handling (pandas-2.0, numpy-1.25), statistical hypothesis testing (scipy-1.11), and planar geometry operations (shapely-2.0).

### Data availability

The mass spectrometry proteomics data have been deposited to the ProteomeXchange Consortium via the PRIDE partner repository with the dataset identifier PXD045356 StA structures were deposited to the EMDB with accession numbers: EMD-18556 (V-ATPase), EMD-18557 (V-ATPase transmembrane partner), EMD-18572 (a segment of CCV isolated from mouse brain tissue), EMD-18574 (a segment of a clathrin-covered endosome isolated from mouse brain tissue), EMD-18578 (empty clathrin basket from mouse brain), EMD-18584 (clathrin triskelion from primary hippocampal neuron culture), EMD-18568 (clathrin triskelion from mouse brain tissue), EMD-18582 (empty clathrin basket from primary hippocampal neuron culture), EMD-18583 (a segment of a CCV from primary hippocampal neuron culture). The structures will be released upon acceptance of the manuscript.

## Acknowledgments

We thank the Core Facility for cryo-Electron Microscopy (CFcryoEM) of the Charité-Universitätsmedizin Berlin for support in the acquisition (and analysis) of the data. The CFcryoEM was supported by the German Research Foundation (DFG) through grant No. INST 335/588-1 FUGG Titan Krios 300 keV Kryo-Transmissions-Elektronenmikroskop. We thank Dr. Thiemo Sprink, Dr. Christoph Diebolder, and Metaxia Stavurolaki for their help with grid preparation and data collection. We thank Dr. Magdalena Schacherl for help with cryo sample preparation and Heike Lerch for help with cell cultivation. We thank Dr. Oleksiy Kovtun, Dr. Natalya Leneva, Dr. Tolga Soykan, Dr. Max Gemmer, Prof. Dr. Reinhard Jahn, and Prof. Dr. Volker Hauke, Dr. Marion Weber-Boyvat, Leonard Roth and Christian Hänig for useful discussions. We thank the Kudryashev Group members Dr. Giulia Glorani, Xiaofeng Chu, and Vasilii Mikirtumov for useful discussions and advice on data processing. We thank Xiaofeng Chu, Daniel Leopoldus, Sabrina Golusik, Elzbieta Wator, and Elena Vazquez Sarandeses for expert annotations of long intravesicular proteusions. We gratefully acknowledge the financial support with a Kekulé fellowship from the Chemical Industry Fund of the German Chemical Industry Association for U.K. The authors thank the Helmholtz Society for funding, M.K. is supported by the Heisenberg Award from the DFG (KU3221/3-1), J.K. is supported by a Walter Benjamin Position from the DFG (458275811) and a fellowship of the DiGiTal program (Berliner Chancengleichheitsprogramm, BCP), C.R. is supported by a Reinhard Koselleck project (399894546) and a NeuroNex project (436260754) from the DFG. M.R. and F.L. are supported by the European Research Council (ERC) Starting Grant (ERC-STG No. 949184). F.L. is supported by Leibniz-Wettbewerb P70/2018.

## Author Contributions

M.K. designed the project. U.K. performed cryo-ET experiments on isolated SVs, subtomogram averaging, and analyzed data. M.R. isolated SVs from mouse brain tissue and performed mass-spectrometry and proteinase K assay. J.K. cultured primary hippocampal neurons on grids and performed sample preparation and cryo-ET data collection on them. A.Y. contributed to the data analysis and simulation of V-ATPases arrangement, M.Z. designed the LuTHy assays with E.W. and performed the experiments. J.R. contributed to the characterization of V-ATPases arrangement. R.L. contributed to the structural analysis of V-ATPases. U.K. wrote the manuscript with input from M.K., J.K., M.R., and A.Y.. M.K., F.L. and C.R. supervised the project, M.K., E.W., F.L., C.R., U.K., and J.K. obtained funding.

