## Supplementary Material for "Molecular architecture of synaptic vesicles"

1. In situ Structural Biology, Max Delbrück Center for Molecular Medicine in the Helmholtz Association (MDC), Berlin, Germany.
2.Depa rtment of Biology, Humboldt University of Berlin, Germany.
3. Leibniz Research Institute for Molecular Pharmacology, Berlin, Germany.
4. Structural Biology of Membrane-Associated Processes, Max Delbrück Center for Molecular Medicine in the Helmholtz Association (MDC), Berlin, Germany
5. Institute of Chemistry and Biochemistry, Freie Universität Berlin, Germany
6.Institute of Neurophysiology, Charité-Universitätsmedizin Berlin, Germany
7. Department of Physics, Humboldt University of Berlin, Germany.
8. Neuroproteomics, Max Delbrück Center for Molecular Medicine in the Helmholtz Association (MDC), Berlin, Germany.
9. Institute of Pharmacy, Freie Universität Berlin, Germany
10. Charité-Universitätsmedizin Berlin, Germany.
11. Institute of Medical Physics and Biophysics, Charité-Universitätsmedizin Berlin, Germany.
12. Indicates equal contribution.

**This PDF file includes:**

Figures S1 to S8

Table S1 to S5

References for Supplementary Material

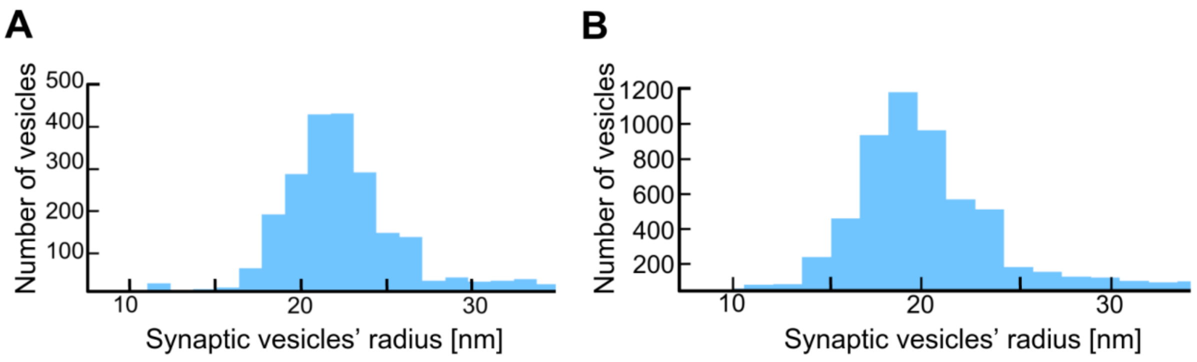


**Figure S1. Histograms representing radii distribution of SVs.** A) in primary hippocampal neurons grown on EM grids B) in isolated SV preparation.

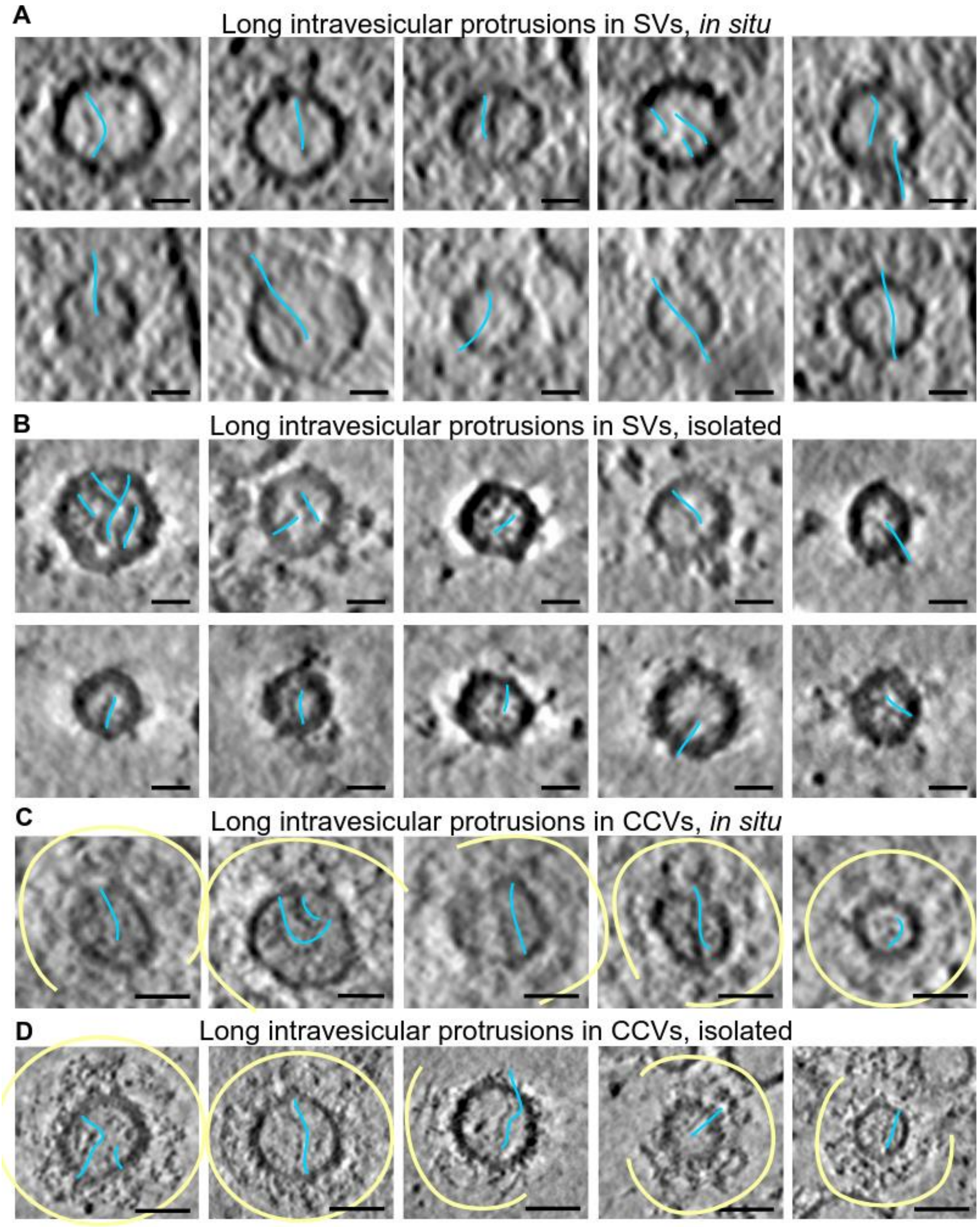


**Figure S2. Examples of long intravesicular densities in SVs and CCVs.** Slices through tomograms of (A) *in situ* SVs (primary hippocampal neurons) (B) isolated SVs (C) *in situ* CCVs (primary hippocampal neurons) (D) isolated CCVs showing long inner protrusions (marked with blue curves next to the protrusion densities). Scale bars (A-B): 25 nm, (C-D): 50 nm. Clathrin cages are shown in yellow (C), (D).

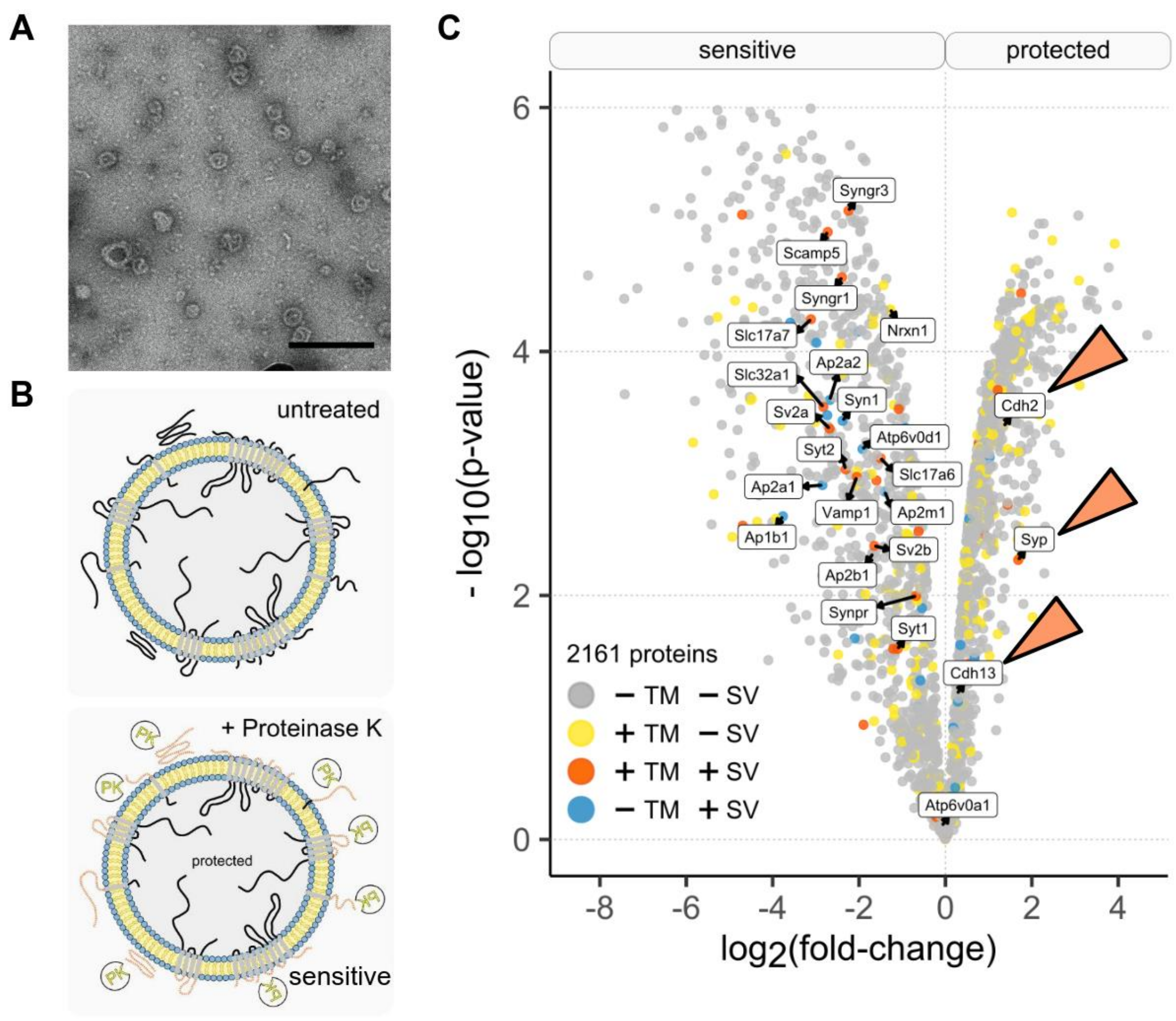


**Figure S3. Mass spectrometry and PK-analysis on SVs.** A) Negative staining performed on the isolated SV sample, scale bar: 200 nm. B) Schematic representation of enzymatic hydrolysis of surface accessible protein domains on isolated murine SVs upon PK treatment. C) Label-free quantitative analysis of PK-treated SVs. Volcano plot with log2 fold-change and p-value of 2161 proteins from three replicates shows proteins susceptible to (negative fold-change) and protected from (positive fold-change) PK treatment. Known protein localization within synaptic vesicles (+SV) and DeepTMHMM-predicted transmembrane domains (+TM) are highlighted. PK-protected proteins are pointed with orange arrows.

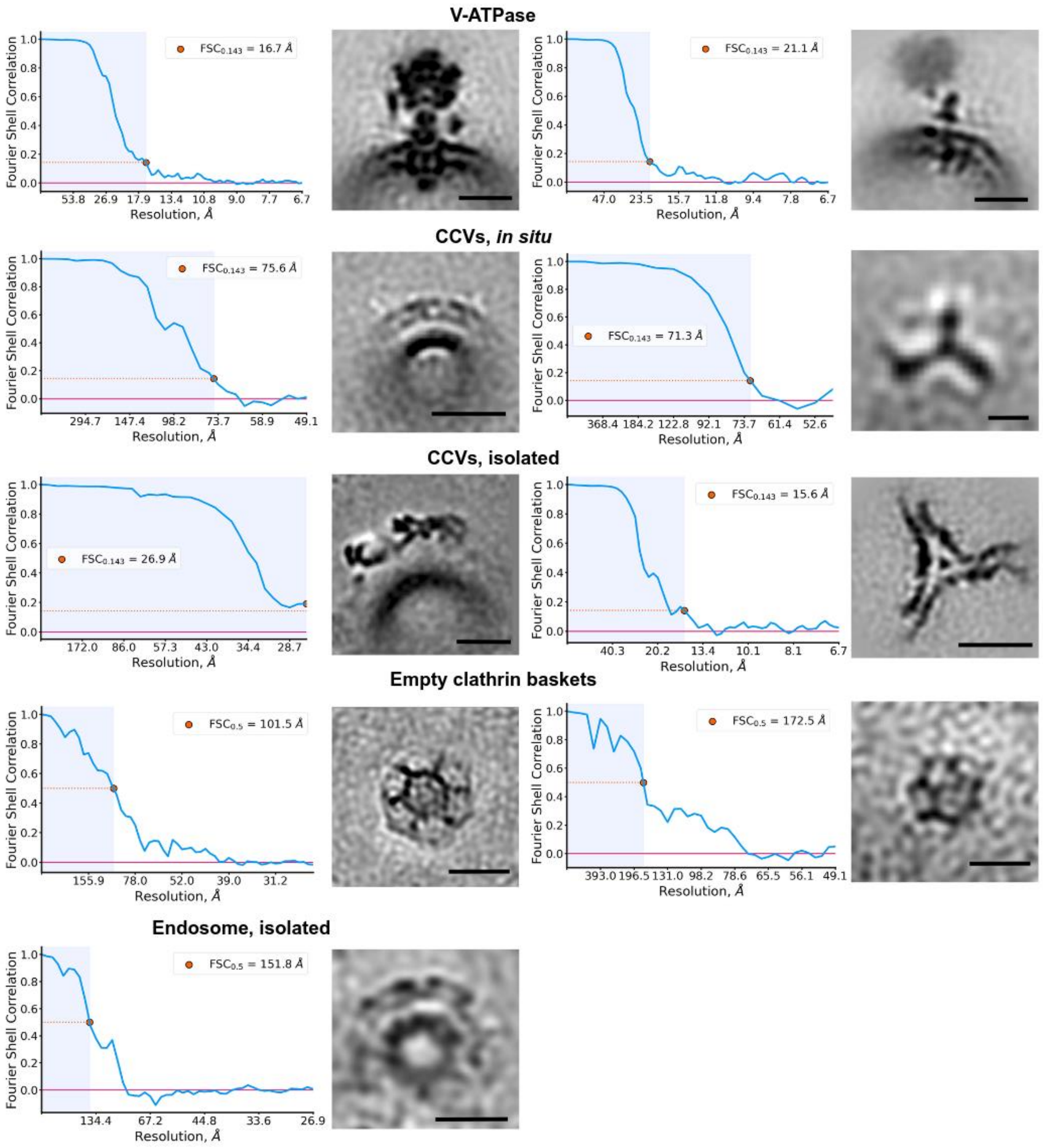


**Figure S4. Structures and resolution estimates of the molecules from the main text.** Fourier Shell Correlations (FSC) curves for the in-text described StA structures. For independent half-set refinements - a low threshold of FSC=0.143 is set up. For empty clathrin baskets and an endosome due to the low number of particles, we could only perform refinement of non-independent half-sets and used a higher threshold of FSC=0.5 for resolution measurement. Scale bars: (V-ATPase) both - 10 nm; (CCVs, *in situ*) left - 50 nm, right - 15 nm; (CCVs, isolated) left - 25 nm, right - 15 nm; (Empty clathrin baskets) both - 50 nm; (Endosome) 50 nm.

A

| Parameters | Syp Q62277 | Synpr Q8BGN8 | Syngr1 O55100 | Syngr3 Q8R191 | Sypl1 O09117 |
|---|---|---|---|---|---|
| Abundance per SV[1] | 31.5 | NA | 2 | NA | NA |
| Presence in MS data[1] | + | NA | + | NA | NA |
| Presence in MS data[2] | + | + | + | + | NA |
| Presence in our MS data | + | + | + | + | - |
| Proteinase K resistance | + | - | - | - | NA |
| Interaction with V-ATPase according to the literature | +[2,3] | -[2] | NA | NA | NA |

B

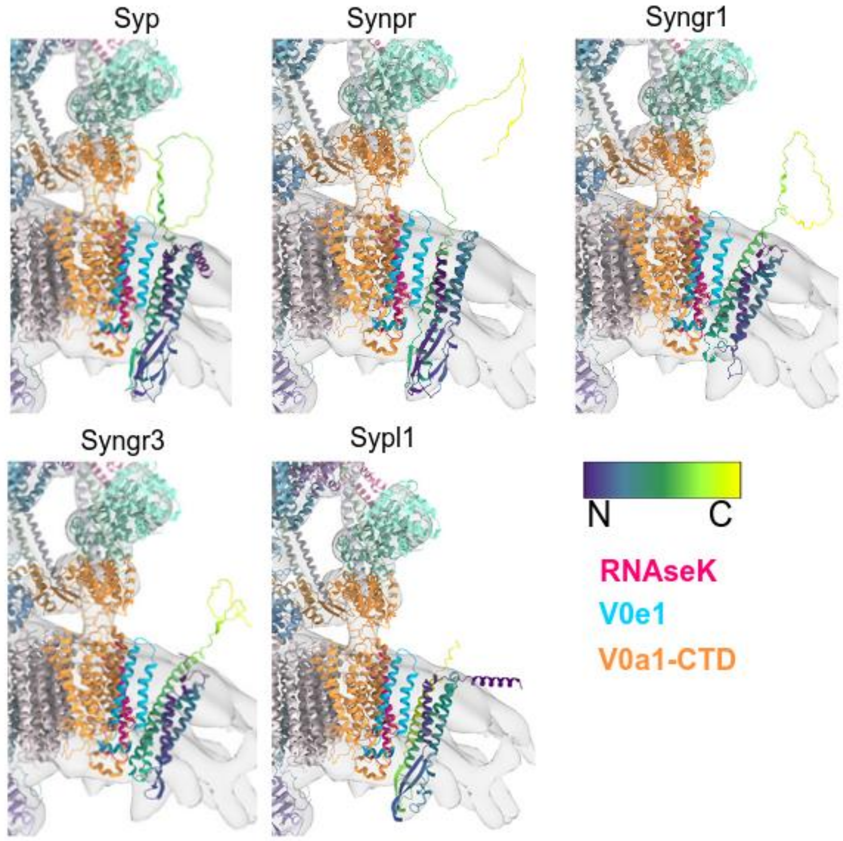


**Figure S5. Candidates for a protein density found in proximity to V-ATPase.** A) A table comparing candidates according to the listed parameters. Among used references: MS data[1,2], immunoprecipitation data[3].B) A rigid body fit of AlphaFold[4]-predicted candidate models to the resolved density next to the V-ATPase Vo domain. V-ATPase model: 6wm2 (PDB)[5]. Candidates are colored in a purple-to-yellow gradient, where purple corresponds to the N terminus and yellow - to the C-terminus.

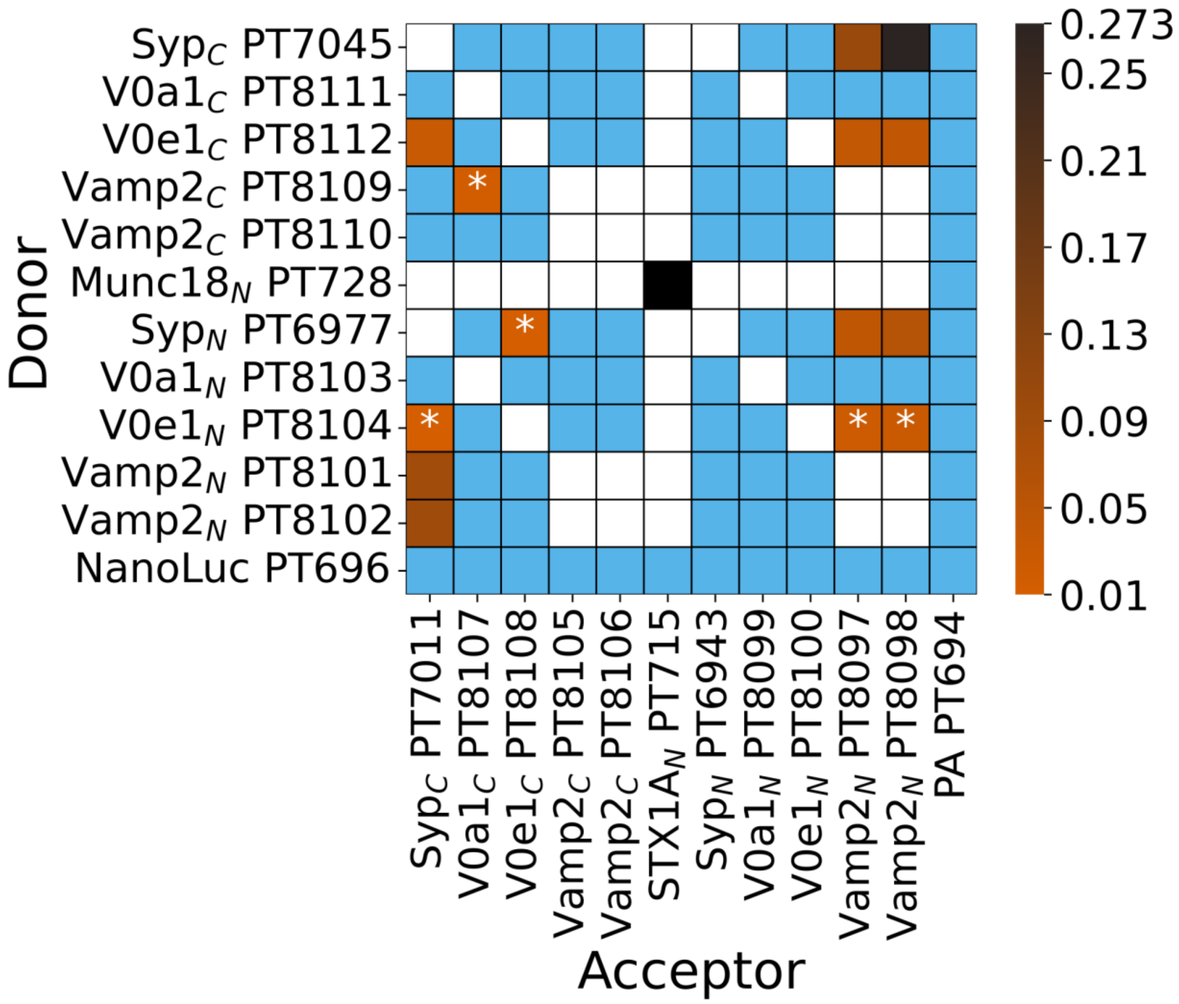


**Figure S6. LuTHy assay performed on V-ATPase Vo e1, V-ATPase Vo a1 domains, Syp and VAMP2.** Control-corrected BRET (cBRET) ratios for the tested pairs of interacting partners as the heatmap, representing protein interaction strength with a color gradient from orange to black. The interactions with observed cBRET values below the cBRET cutoff of 0.01 (cBRET cutoff for cytoplasmic/nuclear/membrane proteins) are represented with blue color. The asterisks (*) annotations represent interactions with cBRET values ≥0.01 (the cutoff for cytoplasmic/nuclear/membrane proteins[6]) but <0.03 (the cutoff for membrane proteins only[6]). The strength of a positive control pair (Munc18-N / STXN1A-N) is shown with a dark black color (cBRET = 0.546), while negative control pairs (NanoLuc-to-all, PA-to-all) are represented with blue color as well. Non-tested pairs are shown by non-colored/white cells.

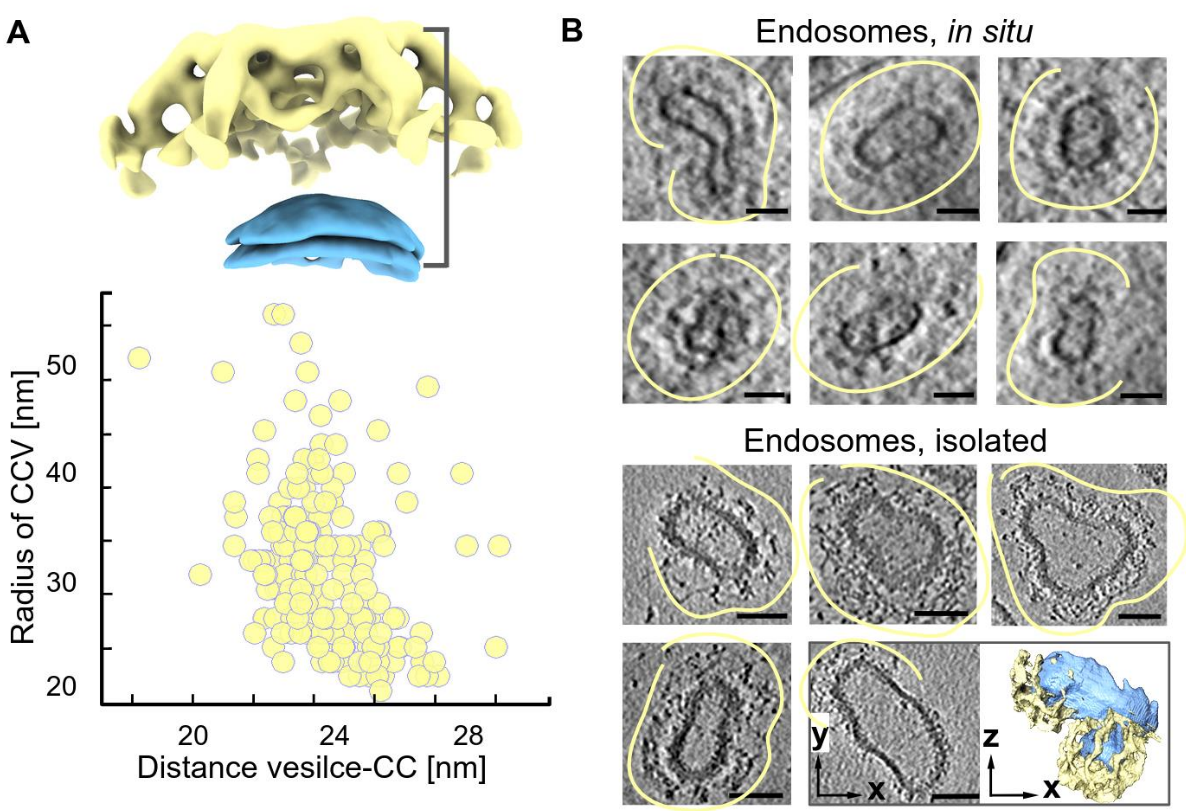


**Figure S7. Clathrin coats (CC), CC-to-coated vesicle distance, and endosomes found *in situ* and the preparation.** A) Top: A volume rendering (ChimeraX) of a CCV surface region, showing CC facet (top, yellow) and CC-enclosed vesicle surface (bottom, blue). The annotation line represents the distance definition for the vesicle-CC measurements. Bottom: Scatter plot of CCVs radii (CC-enclosed vesicle center-to-membrane distances) and the vesicle membrane-to-CC distances. B) Top: Endosomes, observed in neurons. Bottom: Endosomes, isolated from mouse brains, and the AMIRA (Thermo Fisher Scientific) segmentation of one of them. Scale bar is 50 nm.

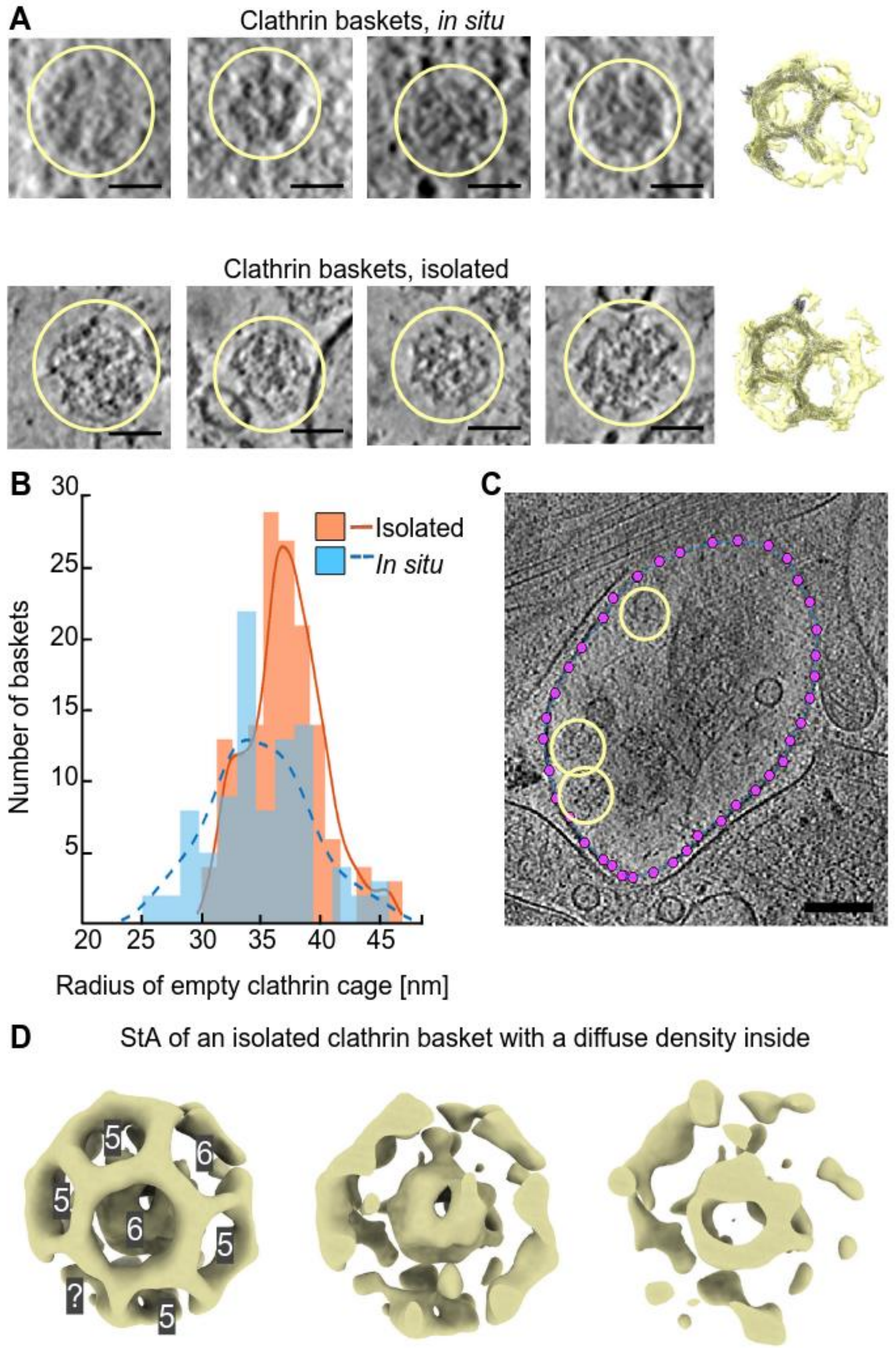


**Figure S8. Visualizations of non-vesicle-containing clathrin baskets.** A) Slices through tomograms of single non-vesicle-containing clathrin baskets *in situ* (upper row) and in isolated fractions (lower row) and their StA structures with the fitted atomic models of the clathrin triskelions (PDB: 6SCT[7]) forming a pentagonal facet. The fitting is performed in ChimeraX. Scale bars: 50 nm. B) Distribution of non-vesicle-coating clathrin baskets radii. C) In-cell clathrin vesicle localization analysis: creating a membrane model using *dynamo_catalogue*, scale bar: 125 nm. D) ChimeraX representation of an isolated basket StA without a vesicle inside. Pentagones or hexagons are shown with numbers 5 or 6 respectively. Three slices show a diffuse density inside.

### Table S1. Data collection and processing statistics for the StA.

**Table S1 Data Processing Statistics**

| | V-ATPase | V-ATPase neighbor | CCVs, *in situ* | clathrin triskelion, *in situ* | clathrin baskets, *in situ* |
|---|---|---|---|---|---|
| **Data collection and processing** | | | | | |
| Microscope | Titan Krios G3i | Titan Krios G3i | Titan Krios G3i | Titan Krios G3i | Titan Krios G3i |
| Magnification | 53,000 x | 53,000 x | 15,000 x | 15,000 x | 15,000 x |
| Voltage (kV), Cs (mm) | 300 kV, 2.7 mm | 300 kV, 2.7 mm | 300 kV, 2.7 mm | 300 kV, 2.7 mm | 300 kV, 2.7 mm |
| Total electron dose ($e^-/Å^2$) | 128…271 | 128…271 | 107 | 107 | 107 |
| Defocus range (µm) | -3.5 to -5.5 | -3.5 to -5.5 | -3.0 to -6.0 | -3.0 to -6.0 | -3.0 to -6.0 |
| Camera | Gatan K3 DED + BioQuantum GIF | Gatan K3 DED + BioQuantum GIF | Gatan K3 DED + BioQuantum GIF | Gatan K3 DED + BioQuantum GIF | Gatan K3 DED + BioQuantum GIF |
| Pixel size (Å) | 3.36 | 3.36 | 24.56 | 24.56 | 24.56 |
| Number of tomograms | 719 | 719 | 34 | 34 | 34 |
| Symmetry imposed | C1 | C1 | C1 | C1 | C1 |
| Initial number of particles | 10204 | 10204 | 15789 | 9384 | 92 |
| Final number of particles | 5361 | 5361 | 1564 | 3658 | 34 |
| Refinement method | Independent half-sets | Independent half-sets | Independent half-sets | Independent half-sets | Non-inependent half-sets |
| Map resolution (Å) | 16.7 | 21.1 | 75.6 | 71.3 | 172.5 |
| FSC threshold | 0.143 | 0.143 | 0.143 | 0.143 | 0.5 |

**Table S1 Data Processing Statistics**

| | CCVs, isolated | clathrin triskelion, isolated | clathrin baskets, isolated | endosomes, isolated |
|---|---|---|---|---|
| **Data collection and processing** | | | | |
| Microscope | Titan Krios G3i | Titan Krios G3i | Titan Krios G3i | Titan Krios G3i |
| Magnification | 53,000 x | 53,000 x | 53,000 x | 53,000 x |
| Voltage (kV), Cs (mm) | 300 kV, 2.7 mm | 300 kV, 2.7 mm | 300 kV, 2.7 mm | 300 kV, 2.7 mm |
| Total electron dose ($e^-/Å^2$) | 128…271 | 128…271 | 128…271 | 128…271 |
| Defocus range (µm) | -3.5 to -5.5 | -3.5 to -5.5 | -3.5 to -5.5 | -3.5 to -5.5 |
| Camera | Gatan K3 DED + BioQuantum GIF | Gatan K3 DED + BioQuantum GIF | Gatan K3 DED + BioQuantum GIF | Gatan K3 DED + BioQuantum GIF |
| Pixel size (Å) | 13.44 | 3.36 | 13.44 | 13.44 |
| Number of tomograms | 147 | 147 | 147 | 5 |
| Symmetry imposed | C1 | C1 | C1 | C1 |
| Initial number of particles | 27498 | 9554 | 265 | 1073 |
| Final number of particles | 1702 | 8006 | 51 | 27 |
| Refinement method | Independent half-sets | Independent half-sets | Non-inependent half-sets | Non-inependent half-sets |
| Map resolution (Å) | 26.9 | 15.6 | 101.5 | 151.8 |
| FSC threshold | 0.143 | 0.143 | 0.5 | 0.5 |

### Table S2. Candidates for the class 1 density - small extra-vesicular domains.

| Candidate class 1 | num per SV in reference[1] | num of TM helixes | TM domain AA | Protein function | UniProt ID | Prediction |
|---|---|---|---|---|---|---|
| Synaptotagmin-1 | | 1 | 58-79 | A component of Ca2+ sensor, which triggers the synchronous release of neurotransmitters in synapses[8]. | P46096 | 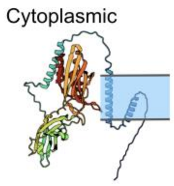 |
| Synaptotagmin-2 | Synaptotagmins - 15.2 | 1 | 61-87 | Inositol-1,3,4,5-tetrakisphosphate (IP4) or inositol polyphosphate-binding protein, potentially involved in synaptic function[9]. | P46097 | 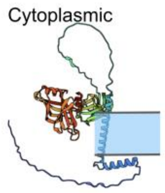 |
| Synaptotagmin-3 | | 1 | 55-75 | Presynaptic Ca2+ sensor, which forces vesicle refilling and short-term synaptic plasticity[10]. | O35681 | 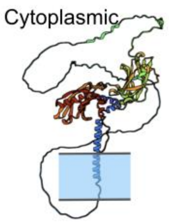 |
| Rab3A | 10.3 | 0 | - | A GTP-binding protein, which is involved in synaptic vesicle transportation to the active zone and docking to the membrane[11]. | P63011 | 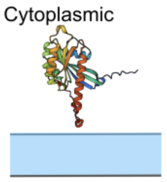 |
| Synapsin-1 | Synapsins - 8.3 | 0 | - | Regulation of axonogenesis and synaptogenesis[12]. | O88935 | 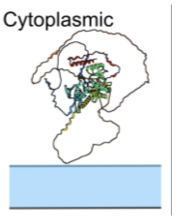 |
| Synapsin-2 | | 0 | - | Desynchronizes γ-aminobutyric acid release in a Ca2+-dependent manner by interaction with presynaptic Ca2+ channels[13]. | Q64332 | 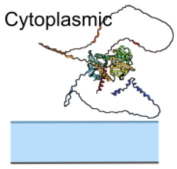 |

| | | | | | | |
|---|---|---|---|---|---|---|
| RalA | - | - | - | A GTP sensor for the GTP-dependent dense core vesicle exocytosis.<br>Not essential for the general secretory pathways[14]. | P63321 | 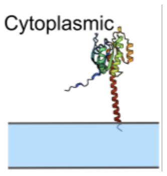 |
| Vesicle-trafficking protein SEC22b | - | 1 | 195-215 | A component of SNARE proteins. Participates in vesicular transport between the ER and the Golgi apparatus[15]. | O08547 | 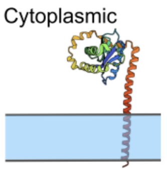 |
| SCAMP1 | 0.8 | 4 | 156-176<br>182-202<br>219-239<br>262-282 | Interacts with EH domain proteins and might participate in endocytosis by directing the assembly of clathrin coats at the plasma membrane[16]. | Q8K021 | 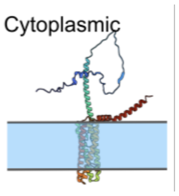 |
| SCAMP3 | - | 4 | 169-189<br>200-220<br>236-256<br>277-297 | Participates in the biogenesis of multivesicular endosomes. Important for EGF receptor sorting into multivesicular endosomes and its targeting to lysosomes by forming intraluminal vesicles within these endosomes *in vitro*[17]. | O35609 | 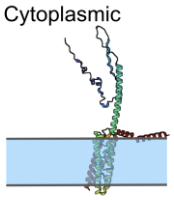 |

(*The N-terminus is shown in blue, and the C-terminus - in red)

#### Table S3. Candidates for the class 2 density - extra- and intra-vesicular domains.

| Candidate class 2 | num per SV in reference[1] | num of TM helixes | TM domain AA | Protein function | UniProt ID | Prediction |
|---|---|---|---|---|---|---|
| SV2a | | 12 | 170-190<br>206-226<br>234-254<br>263-283<br>295-315<br>335-355<br>448-468<br>599-619<br>627-647<br>652-672<br>686-708<br>713-731 | Together with gangliosides, mediates the entry of Botulinum neurotoxin E into neurons[19]. | Q9JIS5 | 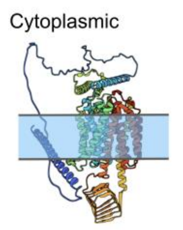 |
| SV2b | SV2 - 1.7 | 12 | 111-131<br>149-169<br>183-203<br>206-226<br>238-258<br>278-298<br>391-411<br>536-556<br>566-586<br>593-613<br>627-649<br>654-672 | Together with gangliosides, mediates the entry of Botulinum neurotoxin E into neurons[19]. | Q8BG39 | 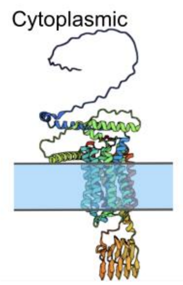 |
| SV2c | | 12 | 155-175<br>192-212<br>227-247<br>249-269<br>281-301<br>321-341<br>438-458<br>579-599<br>610-630<br>637-657<br>671-693<br>698-716 | Mediates a dopamine neuron function; its dysfunction leads to Parkinson's disease which may contribute to dopaminergic dysfunction[20]. | Q69ZS6 | 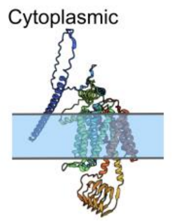 |

(*The N-terminus is shown in blue, and the C-terminus - in red)

### Table S 4. Candidates for the class 3 density - small intra-vesicular domain.

| Candidate class 3 | num per SV in reference[1] | num of TM helixes | TM domain AA | Protein function | UniProt ID | Prediction |
|---|---|---|---|---|---|---|
| Synaptophysin | 31.5 | 4 | 26-49<br>108-131<br>139-162<br>201-224 | Regulates endocytosis during and after neuron stimulation[21]. Regulates synaptic transmission *in vivo.* Redundant and essential in long and short synaptic plasticity. Not required for neurotransmitter release[22]. | Q62277 | 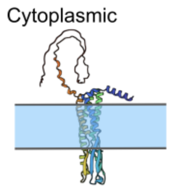 |
| Synaptogyrin-1 | 2 | 4 | 24-44<br>72-92<br>104-124<br>149-169 | Redundant and essential in long and short synaptic plasticity. Not required for neurotransmitter release[22]. | O55100 | 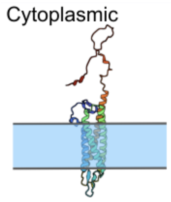 |
| Synaptogyrin-3 | | 4 | 30-50<br>70-90<br>105-125<br>148-168 | Synaptogyrin-3 binding tau protein may cause synaptic dysfunction in progressive supranuclear palsy[23]. | Q8R191 | 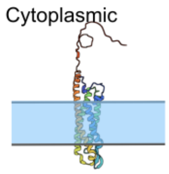 |
| Synaptophysin-like protein 1 | Not available | 4 | 34-54<br>118-138<br>152-172<br>215-235 | Under investigation. | O09117 | 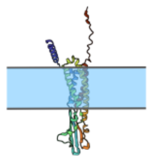 |
| Synaptoporin | | 4 | 5-25<br>82-102<br>115-135<br>179-198 | A putative channel protein of synaptic vesicles[24]. | Q8BGN8 | 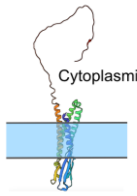 |

(*The N-terminus is shown in blue, and the C-terminus - in red)

**Supplementary Table 5. Candidates for the class 4 density-long intravesicular domain.**

| Candidate class 4 | num per SV in reference[1] | num of TM helixes | TM domain AA | Protein function | UniProt ID | Prediction |
|---|---|---|---|---|---|---|
| Cadherin-13 | Not available | 1 | not annotated | Localized at inhibitory presynapses. CDH13 deficiency in iGABAs increases inhibition and thus decreases excitation/ inhibition balance[25]. | Q9WTR5 | 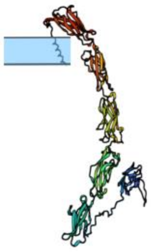 |
| Cadherin-2 | Not available | 1 | 725-745 | Ca2+-dependent homotypic cell adhesion protein[26]. | P15116 | 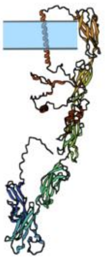 |

(*The N-terminus is shown in blue, and the C-terminus - in red)
